# Vascularized Bioengineered Kidney Using Decellularized Scaffold Recellularized with human Placenta-Derived Angiogenic stem Cells and Kidney Organoids

**DOI:** 10.1101/2025.07.10.664023

**Authors:** Hui Liu, Xi Wen, Yikai Shi, Yijun Zhou, Xinru Chen, Wenpin Song, Xiaoke Zhang, Ying Wang, Junyan Hao, Youyu Ma, Jianse Zhang, Yaoyao Cai, Yongsheng Gong, Feng Lin, Xuguang Du, Junjun Xu

## Abstract

A bioengineered kidney using a decellularized kidney scaffold (DKS) offers a promising solution to the kidney shortage. However, the transplantation of bioengineered kidneys using DKS recellularized with human endothelial, renal cells or others has not yet successfully achieved the vascular and renal reconstruction *in vivo*. In this study, we identified another type of stem cells, designated as human placenta-derived angiogenic stem cells (hPASCs), which serve as seeding cells for the vascularization of DKS. These hPASCs encompass angiogenic subpopulations, exhibit both stem cell properties and the capacity for vascular differentiation. Human fetal kidney organoids (KIO) were established as a source of renal parenchymal cells, comprising renal, immune, and vascular cell populations. We developed a vascularized bioengineered kidney by recellularizing DKS with hPASCs and KIO using a circulation perfusion culture system. The hPASCs bioengineered kidney demonstrated enhanced angiogenesis and reconstructed renal architecture *in vivo*, by transplantation into rat models of renal subcapsular and partial nephrectomy. Furthermore, immunofluorescence and single-cell analyses revealed that the hPASCs revascularized bioengineered kidney regenerated both vascular and renal parenchymal cells within the host. This study offers another strategy for kidney bioengineering and regeneration.

## Introduction

Chronic kidney disease with irreversible loss of renal function will progress to end-stage kidney disease (ESKD)^1^. Patients suffering from ESKD increased from 358, 247 in 2000 to 783, 594 in 2019 with a 41.8% increase^2^. It is estimated that people with ESKD will reach 14.5 million by 2030 while merely 5.4 million will receive kidney replacement therapy (KRT)^3^.However, recipients awaiting transplantation greatly outnumber donors^4^. A bioengineered kidney, utilizing stem cells and biomimetic scaffolds, offers a promising strategy to replace or restore damaged renal function, addressing the shortage of kidney supplies. Adult kidneys consist of over 26 cell types, each with distinct anatomical locations and functions^5^. Selecting suitable cell types and scaffold materials to mimic kidney structures is vital for bioengineered kidneys. However, the lack of an effective vascular network can lead to thrombosis and cell death, hindering *in vivo* transplantation^6^. Constructing an intact vascular network is essential to supply oxygen and nutrients, ensuring graft survival and protecting cells from blood flow shear stress^6, 7^. The source of renal parenchymal cells is another major obstacle in bioengineering kidneys.

Current scaffold materials include synthetic polymers like polycaprolactone, polyvinyl alcohol, and polylactic acid, as well as natural decellularized scaffolds^8^. Synthetic polymers struggle to replicate the *in vivo* microenvironment and native tissue properties, limiting repair capacity due to immunogenicity^9^. In contrast, decellularized kidney scaffolds (DKS) offer improved biocompatibility and lower rejection rates by removing immunogenic components^10^. DKS retains the 3D microstructure of vascular networks, glomeruli, tubules, and signalling molecules, supporting cell adhesion, proliferation, migration, differentiation, and colonization. However, the decellularized ECM can trigger thrombosis after transplantation^11^, and issues like insufficient cell repopulation and inflammatory infiltration lead to ECM reabsorption^12^. Thus, transplanting a DKS alone is insufficient for effective kidney tissue repair and regeneration. The vascular network is crucial for kidney function, supplying oxygen and nutrients to glomeruli and tubules^13^. It comprises vessels of various sizes, structures, and cells, divided into endothelial and supporting types^14^. While human umbilical vein endothelial cells (hUVECs) have been used for vascularization in organs like the kidney^15, 16^, liver^17^, heart^18^, and lung^19^, they fail to develop organ-specific vascular phenotypes and do not effectively connect to host networks^20^. Their poor adhesion also leads to washout, making them suboptimal for bioengineered kidneys due to inefficient vascularization and graft survival issues^21^.To enhance vascularization with hUVECs, co-implantation with mesenchymal stem cells (MSCs) has been explored for bioengineered organs like the kidney^22^ and lung^23^. However, MSCs unable to independently form blood- supporting endothelial tubes^24^, and their differentiation is limited by their tissue origin^25^. Despite advances, significant challenges remain in creating a complete and functional vascular network using hUVECs and MSCs. Thus, the proper source of vascular cells also has been needed to explore for vascularization of bioengineered kidney.

Given the complex renal structure, selecting appropriate renal parenchymal cell types is crucial for regenerating functional bioengineered kidneys. Tissue engineering kidney with DKS recellularized with embryonic stem cell^26^ or kidney cell^15, 27^ has not yet successfully accomplished the reconstruction of vascular and the architecture of renal parenchymal cells *in vivo*. Kidney organoids are miniaturized human-like organs grown from stem or progenitor cells, such as ESCs, iPSCs, or kidney stem cells (KSCs) ^28^. They differentiate into various nephron segments and retain stem cell properties, making them ideal for regenerating renal structures. Adult kidney stem cells can differentiate into nephron progenitor and ureteric bud lineages, forming glomeruli, tubules, and collecting ducts in Matrigel^29^, and are more mature than iPSC-derived cells^30^. No protocol has fully replicated the kidney’s complex structure and function, resulting in organoids lacking immune cells, vascular cells, and other functional structures. Insufficient vascularization limits organoid size (<2 mm) and maturity^31^. After *in vivo* implantation, the integrated ECs are primarily host-derived^32^. Other organoids, like pancreas and brain, co-cultured with ECs, also fail to form host-connected vascular networks^33^. This reliance on host vascularization limits organoid survival and functionality^34^. Whether bioengineering kidney with kidney organoids has accomplished the reconstruction renal parenchymal cell types is unknown.

In this study, we identified another type of stem cells, termed hPASCs, which predominantly consist of subpopulations involved in angiogenesis. These cells exhibit the capacity for vascular differentiation and possess stem cell properties. In 3D Matrigel culture, hPASCs are capable of forming vascular-like tubes comparable to those formed by hUVECs. In the 3D “sandwich” culture model, hPASCs can successfully develop vascular structures, whereas hUCMSCs and hUVECs do not exhibit this capability. Thus, we propose hPASCs as a better seeding cell for vascularizing DKS. KIOs were established as a source of renal parenchymal cells, encompassing populations of renal, immune, and vascular cells. Therefore, we propose a bioengineered kidney constructed using a decellularized scaffold recellularized with human placenta-derived angiogenic stem cells and kidney organoids for vascular and renal structural reconstruction in a rat model.

## Results

### Cultured hPASCs are heterogeneous, angiogenetic potential and multipotent stem cells

The flowchart of Isolation and single cell sequence of hPASCs from full-term placenta was shown in Fig. 1A, a. Single cell RNA sequencing (ScRNA-seq) of the 3rd generation of hPASCs was characterized as functionally enriched 8 clusters which include four angiogenesis, one proliferation and three immunoregulation population (Fig.1B, a-b). Four clusters of hPASCs were enriched in a potent role of angiogenesis functions, including angiogenesis 1 with *ANXA2, SERPINE1*, *ACTG1* and *MYL6*, which were enriched in macrophage cells and angiogenesis^35^; Angiogenesis 2 with *HTRA3*, *SPON2*, *TFPI2* and *AKR1B1* which promote perivascular migration and invade^36–37^; Angiogenesis 3 containing *CXCL1*, *CXCL3*, *CXCL8* and *CCL2*, those were characterized for angiogenic chemokines^38^; Angiogenesis 4 with *STC1, MMP1, MMP14*, and *G0S2*, which were involved invasion and angiogenesis^39–40^.Proliferation including *MT-RNR2*, *MALAT1*, *NEAT1* and *CALD1*, which was related to inhibit cell apoptosis and promote proliferation^41^. Other three clusters of hPASCs were enriched in immunoregulation function: Immunoregulation 1 characterization of innate immune-regulation function for anti-virus which were grouped by the interferon-stimulated genes (*ISG15* and *ISG20*), *OASL*, and *IFIT1*^42, 43, 44^; Immunoregulation 2 included *DIO2* and *COL12A1* which contribute to cell migration^45^;Immunoregulation 3 with *IL1RL1, CCL20* and *SPARC* were related to immune regulation and inflammation^46–47^. Go enrichment of ScRNA-seq data confirmed the molecular characterization of functionally enriched clusters of angiogenesis, proliferation and immunoregulation population (Fig.1B, c).

**Fig. 1.**
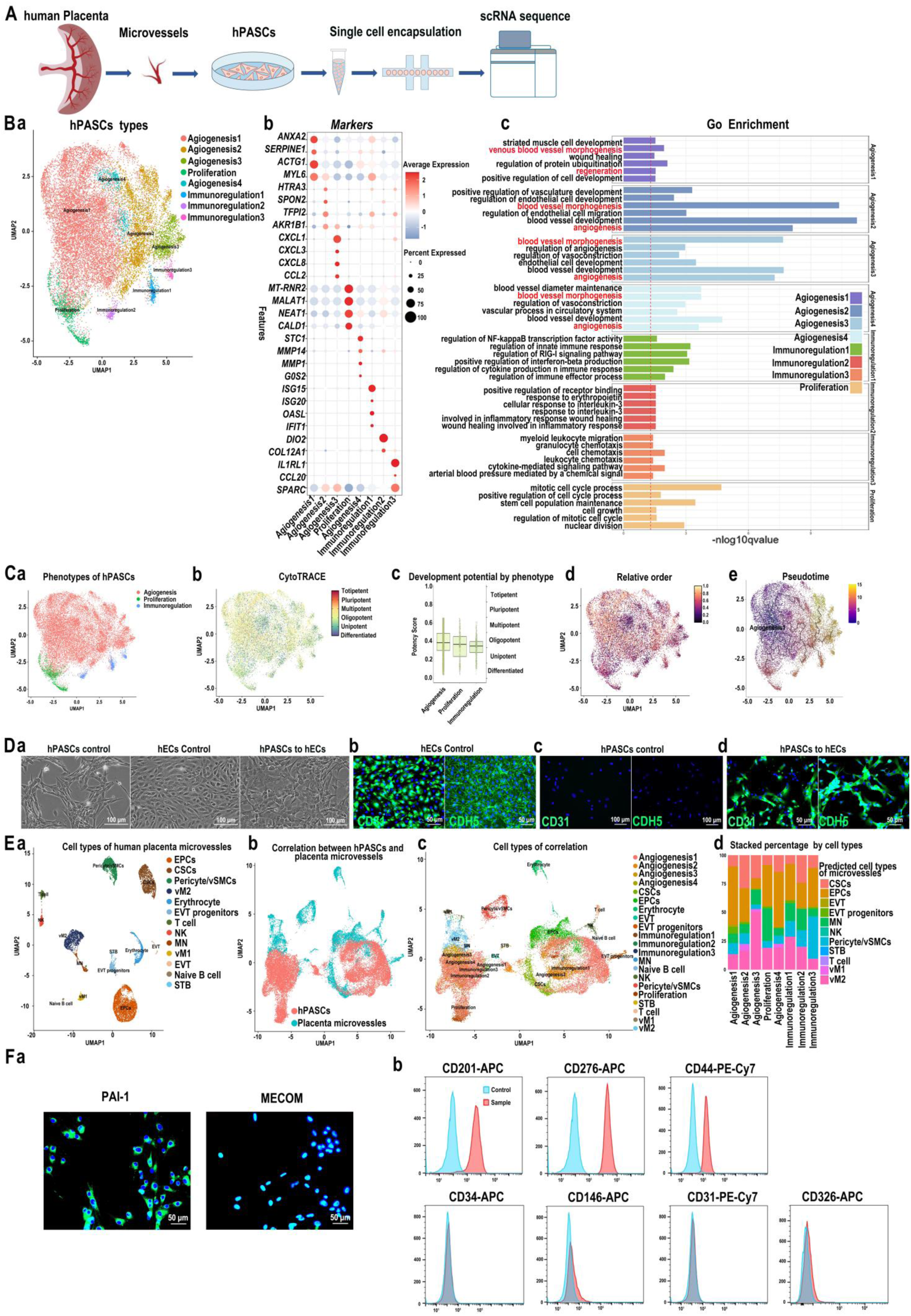
Characterization of hPASCs. (A), The flowchart of isolation of hPASCs and cellular morphology. (B), ScRNA-seq analysis of hPASCs: (a), Cell types enrichment of hPASCs; (b), Markers of hPASCs; (c), KEGG enrichment of ScRNA-seq. (C), Stem cell potentiality of hPASCs: (a), Phenotype; (b), CytoTRACE; (c), Development potential by phenotype; (d), Relative order; (e), Trajectory pseudotime; (D), ECs differentiation of hPASCs: (a), Morphology of hPASCs, hUVECs and ECs differentiation of hPASCs.Scale bar 100 µm; (b), Positive control hUVECs expressing CD31 and CDH5.Scale bar 50 µm; (c), hPASCs were negative for CD31 and CDH5, Scale bar 50 µm; (d), hPASCs were positive for CD31 and CDH5 after ECs differentiation.Scale bar 50 µm. (E), Trace the cellular origins of hPASCs: (a), Cell types of placental microvessels; (b-c) correlation analysis between hPASCs and placental microvessels; (d), Predicts the original cell types of hPASCs by using the placenta microvessles data as a reference dataset. (F), Surface protein identification of hPASCs: (a) Immunofluorescence, hPASCs were positive for PAI-1 and MECOM.Scale bar 50 µm; (b) Flow cytometry.

To validate the stemness of hPASCs, we conducted cellular trajectory reconstruction analysis using gene counts and expression data, which indicated that most hPASCs possess multipotent to unipotent potential (Fig. 1C, a–b). The potential score and relative ordering provided by CytoTRACE further supported the stem cell potential of hPASCs (Fig. 1C, c–d). Pseudotime analysis revealed that the angiogenesis 1 population serves as the initial stage in the developmental trajectory of the clusters (Fig. 1C, e). To confirm the angiogenic potential of hPASCs, we performed in vitro vascular differentiation assays. Morphologically, hPASCs exhibited a polygonal shape (left), which differed from the polygonal paving-stone-like morphology of human umbilical vein endothelial cells (hUVECs) (middle). After endothelial differentiation via EGM-2 medium over three continuous passages, hPASCs changed to the polygonal paving-stone-like morphology (right) (Fig. 1D, a). Immunofluorescence staining demonstrated that control hUVECs were positive for the endothelial markers CD31 and CDH5 (Fig. 1D, b), whereas undifferentiated hPASCs were negative for these markers (Fig. 1D, c). After differentiation, hPASCs exhibited positive staining for CD31 and CDH5 (Fig. 1D, d), indicating they possessed endothelial marker expression during differentiation under these conditions. To assess whether human umbilical cord mesenchymal stem cells (hUCMSCs) possess similar endothelial differentiation capacity, their morphology was observed to shift from polygonal to spindle-shaped after culture in EGM-2 medium (Extended Data Fig.1A, a). However, immunofluorescence staining confirmed that both hUCMSCs and post-differentiation of hUCMSCs remained negative for CD31 and CDH5 ((Extended Data Fig.1A, b–c). Furthermore, both hPASCs and hUCMSCs were cultured in smooth muscle cells (SMCs) differentiation medium. Both cell types expressed the SMCs marker CNN1 (Extended Data Fig.1B, a–b). In summary, these findings demonstrate that hPASCs possess stemness and vascular differentiation capacities, notably differentiating into ECs, but differ from hUCMSCs in their ability to differentiate into ECs.

To trace hPASCs within placental microvessels, scRNA-seq analysis of microvessels was identified 13 cell types: capillary stem cells (CSCs) (marker genes: *PROM1, COL14A1, MMP2, IGFBP3, LUM*), endothelial progenitor stem cells (EPCs) (marker genes: *CD34, PCAM1, SPARCL1, MECOM, KDR, ICAM2*), erythrocytes (marker genes:HBA1, HBG1, HBA2, ALAS2, SNCA), extravillous trophoblasts (EVT) (marker genes: *HLA-G, PAPPA2, PTN, ASCL2*), EVT progenitors (marker genes: *SMAGP, C15orf48, CD24*), mononuclear cells (MN) (marker genes: *BCL2A1, S100A8, G0S2, IL1B, OLR1*), naïve B cells (marker genes: *IGHM, HLA-DRA, CD79A, IGHD, TCL1A*), natural killer (NK) cells (marker genes: *CTSW, NKG7, CLRB1, GZMA, CD247*), pericytes/vascular smooth muscle cells (vSMCs) (marker genes: *MYH11, COX4I2, AREG, GUCY1A2, ITGA1*), syncytiotrophoblasts (STB) (marker genes: *ERVFRD-1, ERVW-1, CYP19A1, KRT23, GDF15*), T cells (marker genes:*ICOS, CAMK4, IL7R, CD3E, CD2*), vascular macrophages 1 (vM1) (marker genes:SPP1, AIF1, C1QA-C, CD68), and vascular macrophages 2 (vM2) (marker genes: *MITF, CD163L1, MRC1, CD163, CCL2, RUNX1*) (Fig. 1E, a; Extended Data Fig.2A, a). The specific marker genes for each cell type are shown in Extended Data Fig.2A, b. GO enrichment analysis further characterized each cell type (Extended Data Fig.2 B). IntegrateLayers function is used to integrate scRNA-seq data from hPASCs and placental microvessles for correlation analysis, aiming to trace the cellular origins of hPASCs. We found that the cell population of hPASCs were closely related to EPCs, CSCs and vM2 of microvessles, while were separated into pericytes/vSMCs and other cell types of placenta microvessles by distance (Fig. 1E, b-c). Furtherly, the MapQuery function predicts the original cell types of hPASCs by using the placenta microvessles data as a reference dataset, revealing that hPASCs predominantly consist of EPCs, CSCs, and vM2 cells (Fig. 1E, d). Trajectory pseudotime analysis between hPASCs and placenta microvessles indicated that EPCs, CSCs and vM2 serves as the developmental origin of hPASCs (Extended Data Fig.2 C).

We aimed to establish a surface molecular panel for the identification of hPASCs. Immunofluorescence analysis demonstrated that hPASCs are positive for the vascular-related marker PAI-1 and the stem cell marker MECOM (Fig. 1F, a). Flow cytometry further revealed that hPASCs express vascular-related stem cell markers, including CD201, CD276, and CD44, while being negative for pericyte marker CD146, hematopoietic stem cell (HSC) and EPCs marker CD34, ECs marker CD31, and epithelial marker CD326 (Fig. 1F, b). These findings suggest that hPASCs possess stem cell properties distinct from pericytes, HSCs, EPCs, and ECs, which have previously been characterized in the human placenta.

In summary, hPASCs comprise angiogenic cell populations, exhibit stem cell characteristics and possess the capacity to vascular cell differentiation. Therefore, we define human placenta-derived angiogenic stem cells as another stem cell type, abbreviated as hPASCs.

### Angiogenic capacity of hPASCs

To validate the vascular-forming capacity of hPASCs, hUVECs and hUCMSCs were used as controls. We conducted in vitro three-dimensional (3D) and 3D “sandwich” culture assays on Matrigel.The model of 3D culture of hPASCs is in Fig.2A, a. Firstly, the hUVECs, hUCMSCs or hPASCs were seeded on Matrigel separately; After 24 hours of culture, hUCMSCs formed spheres, but hUVECs or hPASCs formed vascular-network (Fig. 2A, b). These results demonstrated that hPASCs retained their vascular-forming potential, whereas hUCMSCs did not. Secondly, to evaluate the supportive role of hPASCs in promoting vasculogenesis by hUVECs, co-culture experiments were conducted using hUVECs/hUCMSCs or hUVECs/hPASCs at various ratios (20:1, 10:1, 5:1, 2:1, and 1:1) on 3D Matrigel for 24 and 72 hours, respectively. We observed that hUVECs/hUCMSCs formed spherical aggregates instead of a vascular network structure at any ratios, while hUVECs/hPASCs maintained the vascular network structure at 20:1, 10:1 and 5:1 cell ratio and showed signs of migration at the 2:1 and 1:1 cell ratio (Fig. 2B, a-b), Thus, hPASCs exhibited superior support for vasculogenesis capabilities in comparison to hUCMSCs. Thirdly, in order to further determine migration capacity between hUCMSCs or hPASCs, we labelled hUCMSCs and hPASCs with mCherry, hUVECs with GFP to monitor on 3D culture: After 24 hours culture, hUCMSCs formed spherical aggregates, but hPASCs attached to hUVECs (Fig. 2C, a); After 96 h, hUVECs/hUCMSCs formed spherical aggregates, and neither hUVECs nor hUCMSCs could migrate; hUVECs/hPASCs also formed spherical aggregates, while hUVECs had no migratory ability, but hPASCs migrated on the 3D Matrigel (Fig. 2C, b). Accordingly, Live cell imaging of continuous dynamic showed that hUVECS, hPASCs and hUVECs/hPASCs (1:1) formed vascular network on 3D Matrigel, whereas hUCMSCs or hUVECs/hUCMSCs failed (Fig. 2C, Extended Data Video 1). Lastly, 3D “sandwich” Matrigel culture model was explored to detect vascular-forming potentiality of hPASCs in the three-dimensional space (Fig. 2E, a). 3D “sandwich” culture further showed that hPASCs or hUVECs/hPASCs formed spatial vascular-networks structures, but hUVECs, hUCMSCs and hUVECs/hUCMSCs failed; hPASCs were closely associated with hUVECs, interweaving to form the 3D vascular network (Fig. 2E, b-c). The 3D and 3D “sandwich” models yielded distinct results for hUVECs. This discrepancy is attributed to the fact that the 3D “sandwich” more accurately simulates the spatial structure involved in the complex process of vasculogenesis in the *in vivo* environment of hUVECs.In summary, hPASCs can form vascular network spontaneously, but hUCMSCs and hUVECs are unable to build vascular networks in the 3D “sandwich” model.

**Fig. 2.**
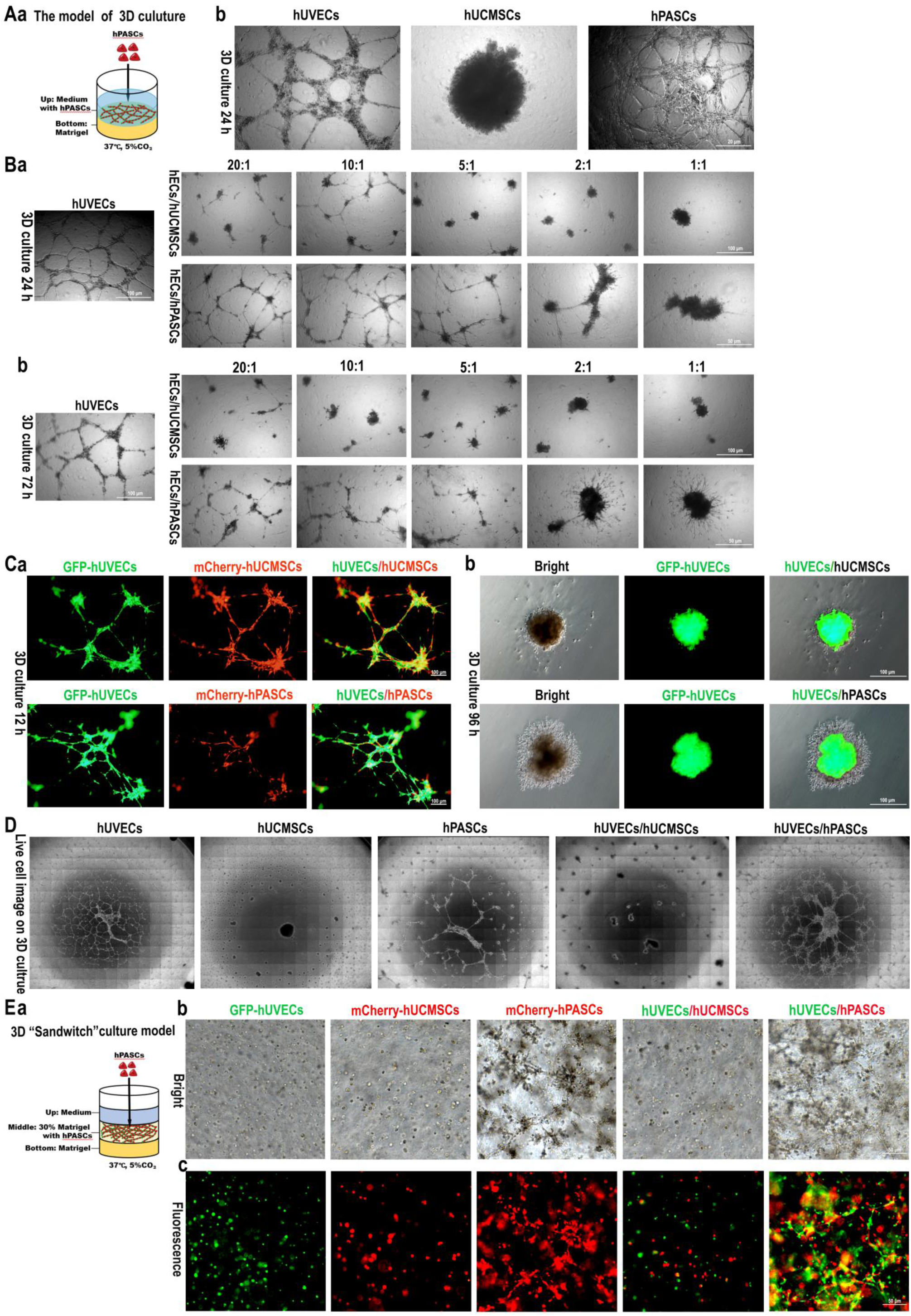
Angiogenic capacity of hPASCs. (A), Vascular-forming potentiality of hPASCs on 3D culture: (a), The model of 3D culture; (b), hPASCs have the vascular-forming potentiality. Scale bar, 20 µm. (B), Different ratio of hUCMSCs or hPASCs with hUVECs were cultured on 3D culture at 24h or 72h.Scale bar, 100 µm (up);50 µm (down). (C), hPASCs interact with hUVECs on 3D culture for 24 hours; (e) Sprouting of hPASCs on 3D culture at 72 h. Scale bar, 100 µm. (D), Live cell imaging of continuous dynamic 3D cultured hPASCs. (E), Angiogenic potentiality of hPASCs on 3D “sandwich” culture: (a), The model of 3D “sandwich” culture; (b-c), hPASCs vascular-network formation on 3D “sandwich” culture at 72h. scale bar, 50 µm.

In summary, on 3D Matrigel culture, hPASCs are capable of forming vascular structures that are highly similar to those formed by hUVECs. However, in 3D “sandwich” models, only hPASCs can generate vascular structures, whereas hUVECs lack this ability. Additionally, hUCMSCs do not exhibit vascular capacity in either 3D or “sandwich” models. These findings indicate that hPASCs possess a unique potential for angiogenesis.

### Isolation and characterization of KIO

The model of isolation, culture, and expansion of the human fetal kidney organoids is shown in Fig. 3A. Two distinct culture mediums of kidney organoids (KIO), PM-hKOCM and D/F12-hKOCM, were developed to culture primary renal cells in a 3D “sandwich” system. The results demonstrated that the PM-hKOCM culture medium maintained the expansion capacity of KIO through sustained proliferation up to passage 6 (P6), while D/F12-hKOCM exhibited a weakened expansion capacity to P3; In contrast, the D/F12-hKOCM system exhibited a progressive decline in proliferative potential, with significantly reduced expansion efficiency observed by passage 3 (P3) and complete loss of proliferative capacity thereafter (Extended Data Fig.3A, a-b). Consequently, PM-hKOCM was selected as the preferred culture system for KIO. The morphology of P3 KIO showed a mixture of folded and cystic shapes (Fig. 3B, a); To assess growth dynamics, P3 KIO were cultured in a 3D “sandwich” system for six days, during which the organoid volume progressively increased, alongside an increase in cell density (Fig. 3B, b). Immunofluorescence staining confirmed the specific marker expression of progenitor cell (PAX2), glomerulus (NPHS1, NPSH2), distal tubule (NKCC2, CDH1), proximal tubule (LTL, LRP2) and vascular cell (EDH3, α-SMA) (Fig. 3B, c). In order to examine whether single cells can large scale regrowth of KIO, we designed the protocol from 3D “sandwich” culture system transferred to 2D culture (Extended Data Fig. 3B, a). Single cells from P3 KIO in a 2D culture with PM-hKOCM were expanded to P6 with cells exhibiting epithelial-like morphology (Extended Data Fig.3B, b). 2D kidney cells expressed PAX2, PODXL, and CDH1 (Extended Data Fig. 3B, c). To investigate the regrowth potentiality of kidney cells in forming KIO, we transfer cells to 3D “sandwich” culture system which showed that the kidney cells from KIO exhibited a regrow kidney organoids (Extended Data Fig.3B, d). Consequently, the PM-hKOCM culture system facilitated the stable passage in 2D culture, and also regrow KIO in 3D “sandwich” culture.

**Fig. 3.**
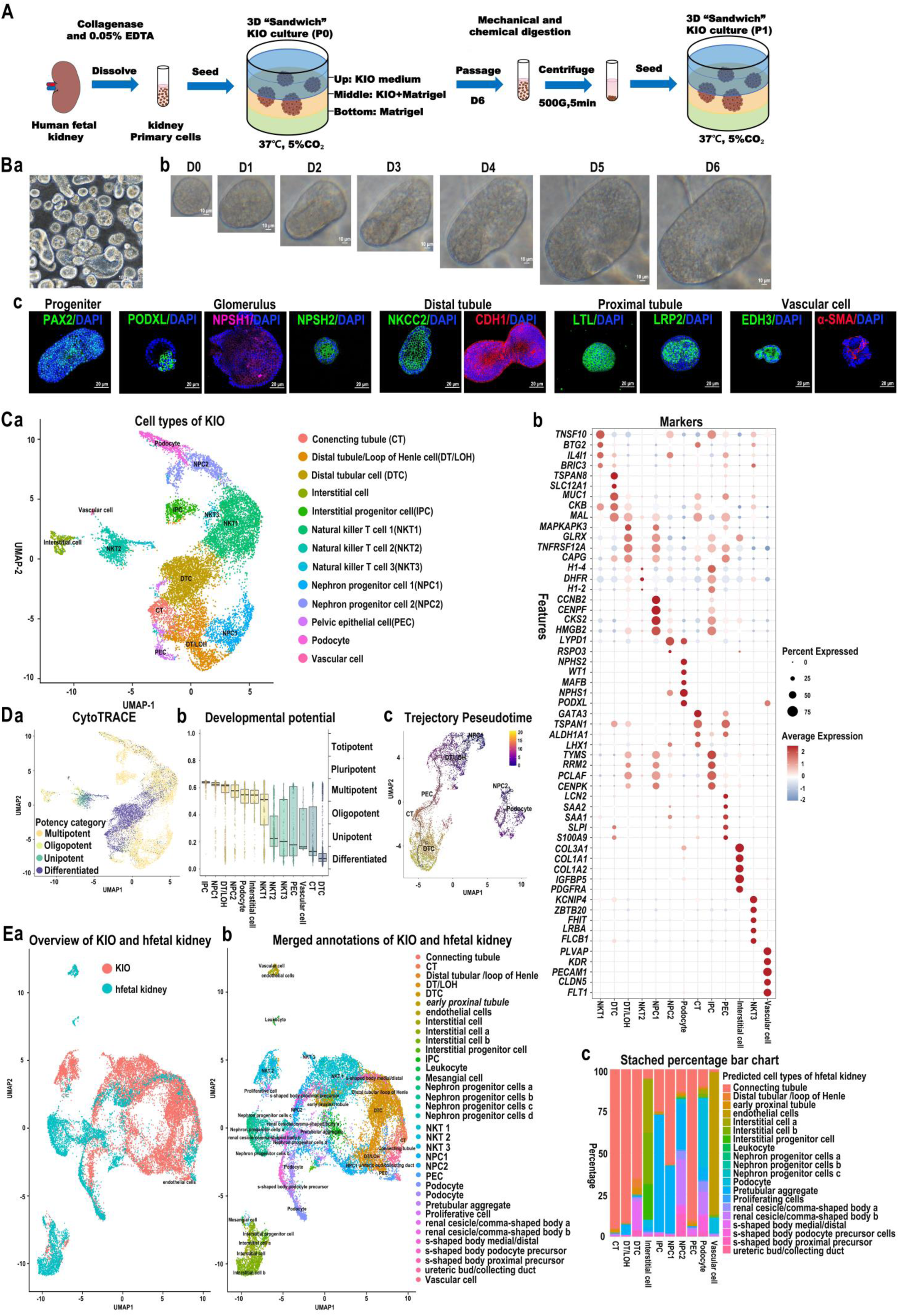
Characterization of KIO. (A), The model of isolation and culture of KIO. (B), The morphology and immunofluorescence of KIO: (a), The morphology of KIO.Scale bar 100 µm; (b), Growth dynamics of KIO.Scale bar 10 µm; (c), Immunofluorescence of KIO.Scale bar 20 µm. (C), ScRNA-seq analysis of KIO: (a) The uniform manifold approximation and projection (UAMP) plot of cell clusters of KIO; (b) Specific marker genes of KIO clusters; (D), (a-b) Analysis of CytoTRACE; (c) Trajectory pseudotime analysis among clusters. (E), Correlation analysis between KIO and human fetal kidney tissue. (a-b) Cellular similarity between organoids and human fetal kidney; (c) Predict the cell types of KIO using human fetal kidney data as a reference dataset.

To elucidate the cellular composition of the kidney organoids, single-cell RNA sequencing (scRNA-seq) was performed on over 10, 000 cells derived from P3 kidney organoids (KIO). This analysis revealed 13 distinct cell types, including nephron progenitors (NPC), distal tubule cells (DTC), distal tubule/loop of Henle (DT/LOH), podocytes, interstitial cells, interstitial progenitor cells (IPC), immune cells, and vascular cells (Fig. 3C, a). Marker genes were identified according to the Cell Marker2.0 dataset (http://www.bio-bigdata.center/index.html). Marker genes expression facilitated the identification of these cell types: nephron progenitors (NPC) were subdivided into NPC1 (characterized by *CCNB2, CENPF, CKS2, HMGB2*) and NPC2 (marked by *LYPD1, RSPO3*), which subsequently differentiate into functional structures such as glomeruli and tubules. Distal tubule cells (DTC) expressed markers such as *TSPANB, SLC12A1, MUC1, CKB,* and *MAL,* confirming their identity. Cells expressing markers associated with the distal tubule/loop of Henle (DT/LOH), including *MAPKAPK3, GLRX, TNFRSF12A,* and *CAPG*, were classified accordingly. Podocytes, identified by the expression of *NPHS2, WT1, MAFB, NPHS1,* and *PODXL,* form the filtration barrier. Connecting tubule (CT) cells expressed *GATA3, TSPAN1, ALDH1A1,* and *LHX1,* indicating their role in urine composition regulation. Renal pelvis epithelial cells (PEC) expressed *LCN2, SAA2, SAA1, SLPI* and *S100A9,* reflecting their involvement in urine storage and transport. Interstitial progenitor cells (IPC) express *TYMS, RRM2, PCLAF,* and *CENPK*, alongside interstitial cells marked by *COL3A1, COL1A1, COL1A2, IGFBP5,* and *PDGFRA*. Vascular cells expressed *PLVAP, KDR, PECAM1, CLDN5,* and *FLT1,* indicating angiogenic potential. Immune cell populations, including subsets of natural killer T (NKT) cells, were identified: NKT1 cells expressed *TNFSF10, BTG2, IL4I1,* and *BIRC3*; NKT2 cells expressed *H1-4, DHFR,* and *H1-2*; NKT3 cells expressed *KCNIP4, ZBTB20, FHIT, LRBA,* and *FLCB1 (*Fig. 3C, b). Overall, the human KIO comprises not only kidney-specific cell types but also immune and vascular components, reflecting a complex and heterogeneous cellular landscape.

CytoTRACE analysis predicted that the subpopulations of IPC, NPC1, DT/LOH, NPC2, podocytes, and interstitial cells retain multipotent capacity, whereas other subpopulations exhibit a more differentiated state (Fig. 3D, a–b).Furthermore, renal parenchymal cells, including DTC, NPC1, NPC2, DT/LOH, CT, podocytes, and PEC, were utilized for analysis to determine the developmental trajectory. Pseudotime trajectory analysis demonstrated that NPC1 differentiates into distal tubule/Loop of Henle (DT/LOH), parietal epithelial cells (PEC), connecting tubules (CT), and distal tubule cells (DTC), while NPC2 differentiates into podocytes, consistent with kidney development (Fig. 3D, c).

The IntegrateLayers function was subsequently employed to integrate scRNA-seq data from KIO and human fetal kidney (GSE114530) for correlation analysis, aiming to assess the cellular similarity between organoids and human fetal kidney. Our analysis revealed an overlap between the two datasets (Fig. 3E, a). Additionally, the cell population composition analysis demonstrated a concordance in vascular, podocyte, interstitial, and other cell types (Fig. 3E, b). Furthermore, the MapQuery function was utilized to predict the cell types of KIO using human fetal kidney data as a reference dataset, indicating that KIO shares similarities with most cell types of the human fetal kidney (Fig. 3E, d). Therefore, the cellular composition of KIO appears capable of recapitulating the majority of cell types found in the human fetal kidney.

### hPASCs vascularized bioengineered kidney construction and filtration function *in vitro*

The procedures for constructing the vascularized bioengineered structure and its transplantation were shown in Fig. 4A. The isolated rat kidney was light yellow after perfusion with 1 ɡ/L sodium heparin solution via the renal artery (Fig. 4B, a), faded after perfusion of 1% (v/v) Triton X-100 solution (Fig. 4B, b) and eventually became white and transparent after perfusion of 8 ɡ/L SDS solution (Fig. 4B, c). The transparent, white, and uniform texture of the DKS allowed a clear vision of the structure (Fig. 4B, d). Scanning electron microscopy (SEM) exhibited the smooth surface of the ECM. The scaffold, which retained a good 3D pattern, had a clear spatial honeycomb structure (Fig. 4B, e). The colour of the kidney scaffold changed from a cell-free milky white (Fig. 4C, a) to pink (Fig. 4C, b), after hPASCs were injected into it via indwelling cannulas connected to the renal artery and vein. a circumfusion culture system was designed for the circulation of medium from the artery to the vein of the bioengineered kidney, which provided sufficient oxygen and nutrient for cell growth (Fig. 4D). When 2 × 10^7^ hPASCs were injected into DKS culture for 4 days, A total of 1×10^6^ KIOs were injected to KDS after 24 hours culture for immunofluorescence detection of hPASCs and KIO location in DKS. hPASCs were labeled with mCherry to facilitate the tracing of their localization within the DKS. Immunofluorescence analysis demonstrated that hPASCs adhered to the scaffold and contributed to the formation of cavity-like structures (Fig. 4E, a). Additionally, NKCC2 immunofluorescence was employed to detect the localization of KIO within the DKS. The results indicated that KIO adhered to the wall of the DKS cavity but did not exhibit the same level of attachment or adhesion as hPASCs (Fig. 4E, b). This discrepancy is likely attributable to the relatively short incubation period, as KIO was only injected 24 hours prior to analysis.

**Fig. 4.**
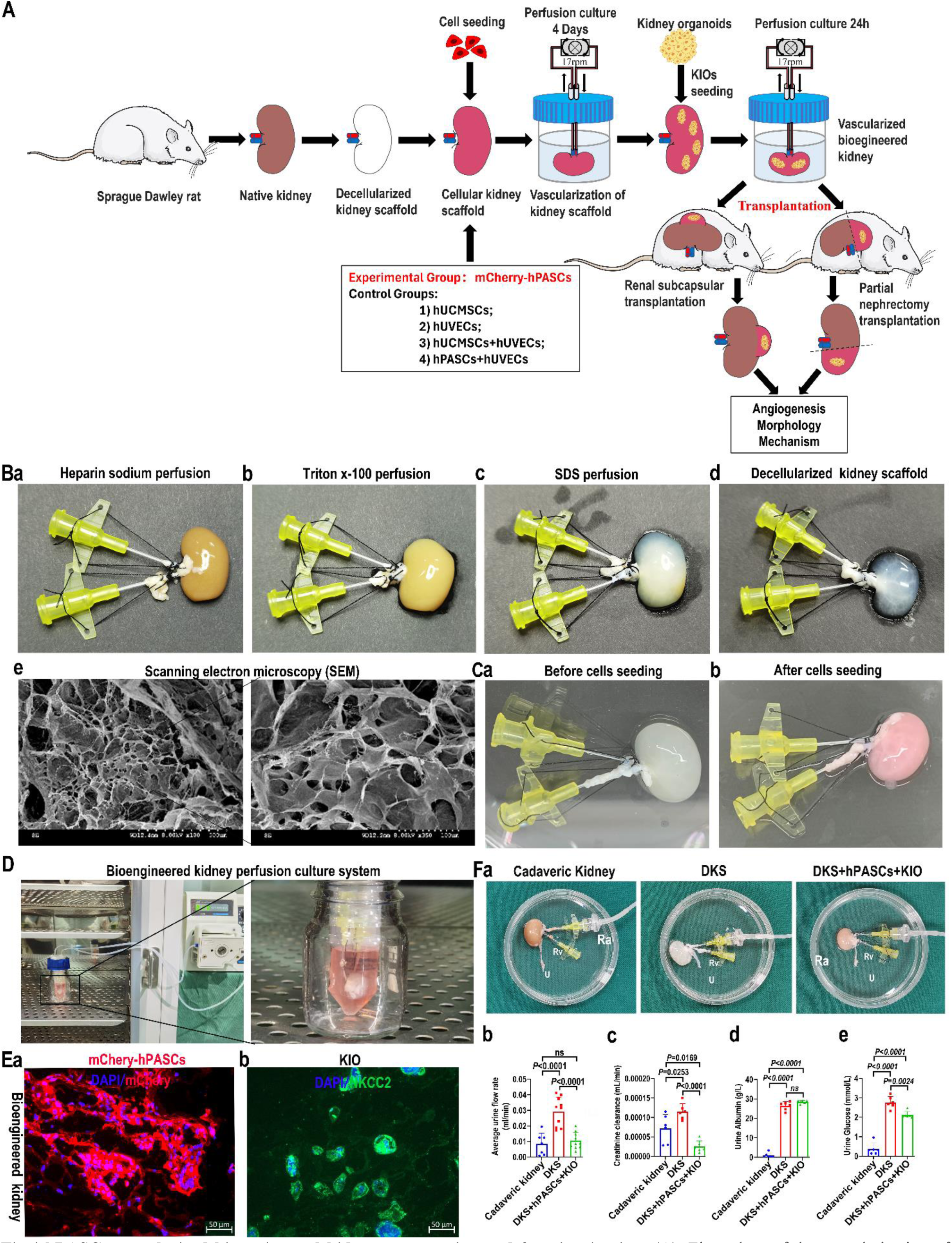
hPASCs vascularized bioengineered kidney construction and function *in vitro*. (A), Flow chart of the vascularization of bioengineered kidney and *in vivo* transplantation. (B), Preparation of DKS: (a), Perfusion with sodium heparin solution; (b) Perfusion with Triton X-100 solution; (c), Perfusion with SDS solution; (d), The photograph of DKS; (e) SEM of ultrastructure of DKS. (C), Recellularization of DKS: (a), Before cell seeding; (b), After cell seeding. (D), Circumfusion culture of bioengineered kidney. (E), Immunofluorescence detection of location of hPASCs and KIO within DKS: (a), hPASCs adhered to the scaffold and contributed to the formation of cavity-like structures; (b), KIO adhered to the wall of the DKS cavity but did not exhibit the same level of attachment or adhesion as hPASCs; Scale bar, 50µm. (F), The filtration function of hPASCs vascularized bioengineered kidney: (a), The scheme of filtration of perfusion fluid; (b), The average urine flow rate (mL/min), *n*=7, 10, *P*<0.05; (c)Creatinine clearance rate, *n*=6, *P*<0.05; (e) Albumin retention rate, *n*=6, *P*<0.05; (d) glucose reabsorption function, *n*=6, *P*<0.05. Ra, renal artery, Rv, renal vein; U, ureter. Statistics analysis was performed using Tukey’s multiple comparisons test.

In order to test the filtration function of hPASCs vascularized bioengineered kidney *in vitro*, perfusion fluid was filtered and reabsorbed through cadaveric kidney, DKS, and DKS+hPASCs+KIO group. The perfusion fluid was filtered through cadaveric kidneys, DKS group, and DKS+hPASCs+KIO group (Fig. 4F, a). The average urine flow rate (mL/min) was calculated for each group. Statistical analysis revealed that the average urine flow rate in DKS group is faster than in the cadaver kidney group (*P*<0.001, *n*=7) and DKS+hPASCs+KIO group *P*<0.001, *n*=10); while there is no difference between the cadaver kidney and DKS+hPASCs+KIO group (Fig. 4F, b; Extended Data table 1).These findings indicate that the DKS+hPASCs+KIO group exhibited an improvement in renal concentrated urine and reabsorption function. Furtherly, the creatinine clearance rate of DKS group is higher than that of the cadaveric kidney group (*P*<0.0253, *n*=6) and DKS+hPASCs+KIO (*P*<0.0001, *n*=6); The reduction in clearance rate can be attributed to the lower urine production because of the presence of an intact filtration membrane (Fig. 4F, c; Extended Data table 2); Thus, DKS+hPASCs+KIO group displayed a significantly creatinine clearance. The DKS and DKS+KIO+hPASCs group had no difference of albumin absorption, but both of them were higher than that of the cadaveric kidney group, indicating that DKS+KIO+hPASCs had no albumin absorption (Fig. 4F, d; Extended Data table 3). The DKS group exhibited a significantly higher level of glucose in the urine compared to the cadaveric kidney group (*P*<0.0001, *n*=6), indicating the loss of glucose reabsorption function in the decellularized kidney; but, the DKS+KIO+hPASCs group showed a partial recovery in glucose reabsorption function when compared to the DKS group (*P*=0.0024, *n*=6) (Fig. 4F, e). Collectively, hPASCs vascularized bioengineered kidney with KIO exhibited a certain filtration function *in vitro*.

### Vascularization of DKS with hPASCs *in vitro*

Before assessing the vascularization of the DKS with hPASCs, we first analysed the biomechanical disparities among hUVECs, hUCMSCs and hPASCs. Traction force microscopy (TFM) measurements revealed differences in the representative traction force (RMS) and the total strain energy (TSE) exerted by individual cells. RMS maps of individual cells (Extended Data Fig.4A, a). RMS traction of hUCMSC is higher than that of hUVECs (*P*<0.0001, *n*=17) and hPASCs (*P*<0.0001, *n*=17). Meanwhile RMS traction of hPASC was higher than that of hUVECs (*P*=0.0074, *n*=17) (Extended Data Fig.4A, b); The total strain energy (pJ) of hUCMSCs were higher than hUVECs (*P*<0.0001, *1*=17) and hPASCs (*P*<0.0001, *n*=17), there was no difference between hPASCs and hUVECs (Extended Data Fig.4A, c; Extended Data table 4). Therefore, hUCMSCs exhibited the highest RMS traction forces and total strain energy (pJ) compared to hUVECs and hPASCs. We hypothesize that hPASCs possess a stronger adhesion capability within the DKS relative to hUCMSCs.

After 2×10^7^ hPASCs were implanted into the DKS and cultured in the perfusion culture system (37℃, 5% CO2) for four days, then1×10^6^ KIOs were injected into it in the same way and co-cultured for 24 hours. Fluorescent labeling of mCherry-hUCMSCs, mCherry-hPASCs, and GFP-hUVECs was used to assess vascular formation. The DKS+hPASCs+KIO and DKS+hUVECs+KIO groups formed CD31-positive vascular structures; The DKS+hPASCs+hUVECs+KIO group showed vascular structures with hUVECs adhering outside hPASCs, indicating preferential scaffold adhesion of hPASCs; Groups DKS+hUCMSCs+KIO and DKS+hUCMSCs+hUVECs+KIO showed cell aggregation without organized vasculature; No vascular structures or CD31 expression were observed in DKS and DKS+KIO groups (Extended Data Fig. 4B, a). Similarly, α-SMA staining confirmed these results, except for DKS+KIO, which expressed α-SMA (Extended Data Fig. 4B, b). Overall, hPASCs can form vascular structures and interact with hUVECs, whereas hUCMSCs do not.

SEM revealed that the scaffold’s spatial structure was well preserved in the DKS+hPASCs+KIO group after continuous perfusion culture, while other groups showed varying degrees of collapse and structural incompleteness; additionally, cell connection structures were observed in the DKS+hPASCs+KIO group (Extended Data Fig. 4C, a).TEM showed that in DKS+hPASCs+KIO group, cells extended pseudopods attaching to the scaffold and connection structures between cells; in other groups, cell debris, vacuoles and apoptotic bodies were observed (Extended Data Fig. 4C, b).

Conclusively, hPASCs possess strong adhesion ability, form vascular structures with KIO within DKS.

### hPASCs vascularized bioengineered kidney promote angiogenesis in renal subcapsular transplantation

Bioengineered kidneys were harvested 14 days after renal subcapsular transplantation (Extended Data Fig. 5A). To determine the angiogenic effect of hPASCs after subcapsular transplantation, vessel density was assesed by fluorescent analysis using the vascular endothelial-specific marker CD31; CD31^+^ vessels in DKS+hPASCs+KIO group was significantly more abundant than that of other groups (Extended Data Fig. 5B, a). Statistical analysis revealed that the CD31^+^ vessels density was significantly higher in the DKS+hPASCs+KIO implants than in DKS (*n*=3(*P*=0.002), DKS+KIO (*n*=3, *P*=0.046), DKS+hUCMSCs+KIO (*n*=5, *P*=0.004) and DKS+hUVECs+KIO (*n*=5, *P*=0.001) implants, but CD31^+^ vessels density in the DKS+hUVECs+KIO, DKS+hUCMSCs+KIO, DKS+KIO, and DKS implants showed no statistical difference (Extended Data Fig. 5B, b; Extended Data table 5), indicating that hPASCs had stronger angiogenic effect compared with hUVECs and hUCMSCs, co-implantation of hUCMSCs or hPASCs with hUVECs did not remarkably improved angiogenesis after subcapsular transplantation.

HE staining demonstrated that the DKS+hPASCs+KIO implants formed typical glomerular-like structures similar to native kidneys (Extended Data Fig. 5C, a-b); the tubular-like structures appeared more immature compared to native tissue (Extended Data Fig. 5D, a-b). Furthermore, fluorescent staining confirmed that implant of DKS+hPASCs+KIO expressed the markers of glomerulus (NPHS1, NPHS2), collecting tube (AQP2), distal tube (NKCC2, CDH1), and proximal tube (LRP2) the native tissue was as control (Fig. 5E, a-b). Thus, the transplantation of hPASCs vascularized bioengineered kidney can reconstruct the kidney structure.

**Fig. 5.**
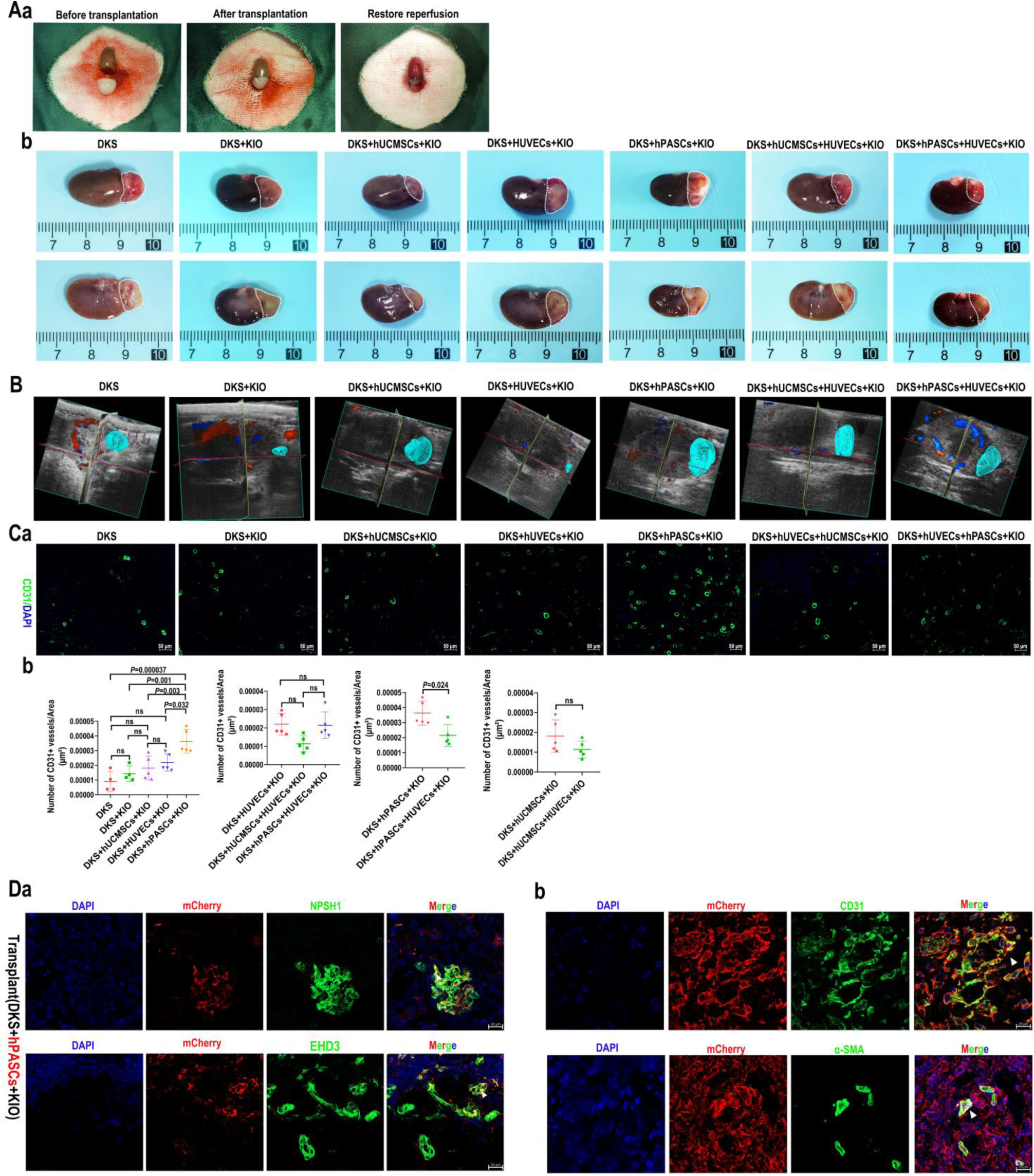
hPASCs vascularized bioengineered kidneys promote angiogenesis in rat partial nephrectomy model. (A), The scheme of transplantation: (a), transplantation of bioengineered kidney; (b)The photograph of bioengineered kidney 14 days post transplantation; (B), Ultrasound 3D vascular modelling in the implant. (C), Angiogenesis of hPASCs: (a), Immunofluorescence of CD31 staining. Scale bar 50 µm; (b), Statistical analysis of the density of vascular structures: DKS and DKS+KIO groups (*n*=4); other groups (*n*=5); *P*<0.05. (D) hPASCs invoved in vascular formation in the host. (a), hPASCs formed the network of blood vessels in glomerulus and co- expression of EHD3; (b), hPASCs were also co-expressed CD31 and α-SMA. For multiple group comparisons, the Kruskal Wallis rank sum test was used when the variances were unequal, and the Bonferroni method was used for multiple comparison. *P*<0.05 was considered statistically significant.

Collectively, hPASCs vascularized bioengineered promote angiogenesis, take part in the vascular and renal structure formation in the renal subcapsular transplantation.

### hPASCs vascularized bioengineered kidneys promote angiogenesis in rat partial nephrectomy model

To validate the role of hPASCs bioengineered kidney in promoting vascularization and renal structural reconstruction in a rat model, a one-third partial nephrectomy model was utilized for transplantation. The caudal part of the rat kidney was resected aseptically, and a bioengineered kidney of similar volume was transplanted; following blood reperfusion, the transplanted bioengineered kidney tissue exhibited a characteristic red coloration (Fig. 6A, a). Kidneys were harvested 14 days post-transplantation. A clear boundary between the implanted tissue and the residual kidney was observed, and all group implants detectable in longitudinal sections (Fig. 6A, b).

**Fig. 6.**
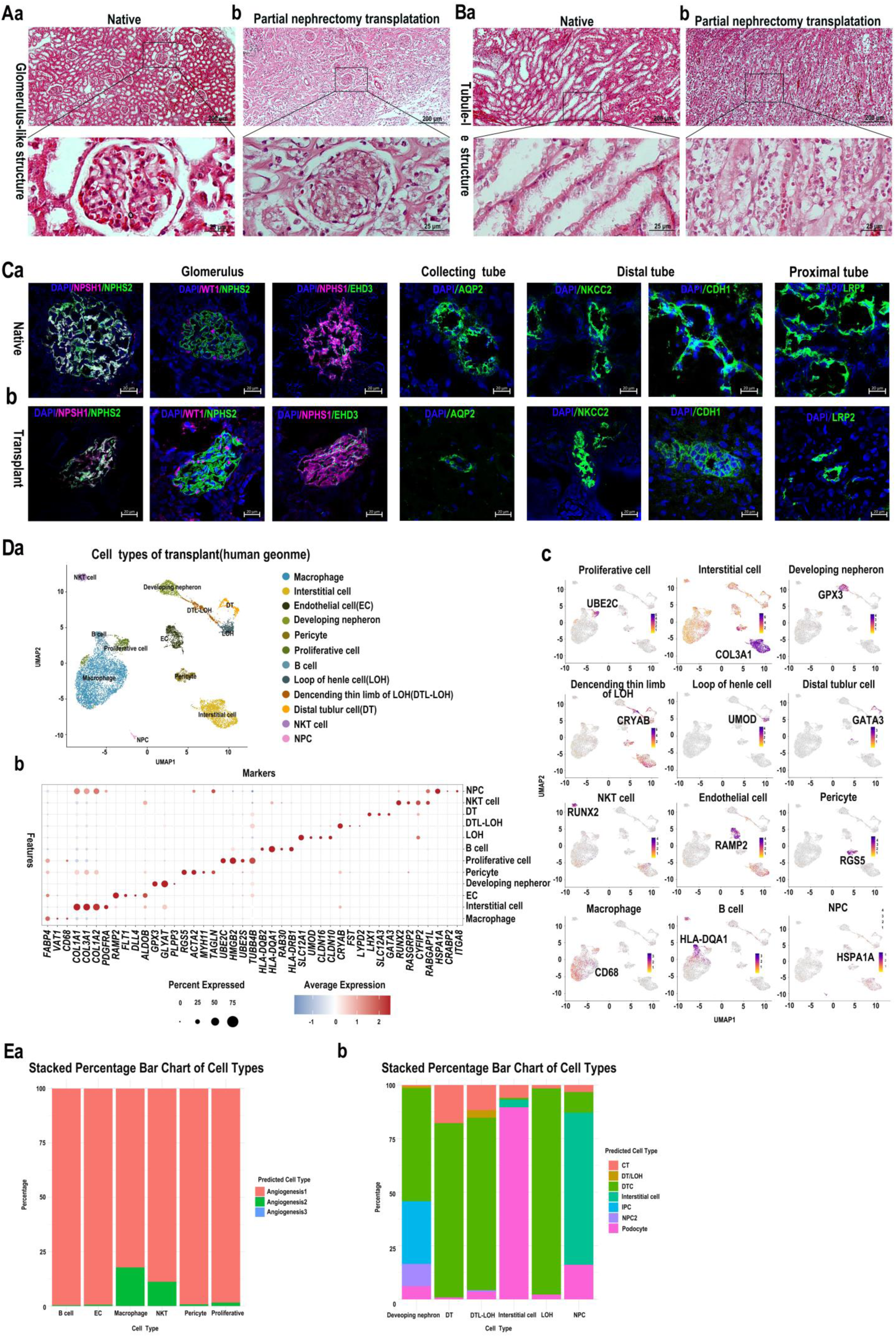
Regeneration of hPASCs vascularized bioengineered kidney in rat partial nephrectomy model: (A), HE staining analysis of Glomerulus-like structure: (a), Native group; (b), DKS+hPASCs+KIO implant group. Scale bar 200 µm (up); 25 µm (down). (B), HE staining analysis of Tubule-like structure: (a), Native group; (b), DKS+hPASCs+KIO implant group.200 µm (up); 25 µm (down). (C), Fluorescent staining of implant: (a), The native tissue was as control; (b), Implant were positive for markers of glomerulus (NPHS1, NPHS2, EHD3), collecting tube (AQP2), distal tube (NKCC2, CDH1), and proximal tube (LRP2). Scale bar 20 µm. (D), ScRNA-seq analysis of implant of DKS+hPASCs+KIO using human genome: (a), Human cell types; (b), Marker genes of each cell type; (c), Specific markers. (E), The MapQuery function predicts the original cell types; (a), Vascular and immune cells were derived from hPASCs; (b), Kidney parenchymal cells originated from KIO.

On day 14 post-partial nephrectomy, each group received a sutured bioengineered kidney implant. Ultrasound 3D vascular modelling revealed that the DKS+hPASCs+KIO group exhibited higher vascular density compared to other groups, with relatively lower densities observed in the DKS+KIO, DKS+hUVECs+KIO, and DKS+hUCMSCs+hUVECs+KIO groups (Fig. 6B). Fluorescent staining demonstrated a greater number of CD31^+^ vessels in the DKS+hPASCs+KIO group (Fig. 6C, a). Statistical analysis confirmed that vascular density in the DKS+hPASCs+KIO group was significantly higher than in DKS+hUVECs+KIO (*n*=5*, P*=0.032), DKS+hUCMSCs+KIO (*n*=5*, P*=0.003), DKS+KIO (*n*=4*, P*=0.001), and DKS alone (*n*=4*, P*=0.000037), no significant differences were observed among the other groups; Additionally, the vascular density in the DKS+hPASCs+KIO group exceeded that of the DKS+hUVECs+hPASCs+KIO group (*n*=5*, P*=0.024) (Fig. 6C, b; Extended Data Table 6). These findings suggest that hPASCs exhibit stronger angiogenic capacity compared to hUCMSCs or hUVECs, and that combined transplantation with hUVECs does not significantly enhance angiogenesis in the host. Fluorescence tracking of hPASCs confirmed that hPASCs formed the network of blood vessels in glomeruli with NPSH1 and mCherry expression, and partially expressed the EDH3 of glomerular endothelial cells (Fig. 6D, a), hPASCs were also co-expressed CD31 and α-SMA (Fig. 6D, b). Thus, these results indicated that hPASCs participate in the vascular formation of glomeruli and vessels in the host.

### hPASCs vascularized bioengineered kidney reconstructed renal structure in rat partial nephrectomy model

Fourteen days after the transplantation of engineered kidneys from the DKS+hPASCs+KIO group into SRG rats following partial nephrectomy, transplant samples were collected for histological, fluorescence, and scRNA-seq analyses.First, cross-sectional HE staining showed that the implants of the DKS+hPASCs+KIO group exhibited glomerular-like structures resembling native glomeruli, which included clear capillary loops and the Bowman’s capsule (Fig.7A, a-b); however, the tubular structure appears more immature compared to the native structure (Fig.7B, a-b). Second, fluorescent staining confirmed that implant of the DKS+hPASCs+KIO expressed the markers of glomerulus (NPHS1, WT1and NPHS2), collecting tube (AQP2), distal tube (NKCC2 and CDH1), and proximal tube (LRP2), the native tissue was as control (Fig.7C, a-b). Thus, the implant of DKS+hPASCs+KIO was not only contributed to the glomerulus, but also tubular formation in the group.

Furtherly, ScRNA-seq analysis of implant of DKS+hPASCs+KIO using human genome, cell types of implants contained the human kidney parenchymal cells (interstitial cells, developing nephrons, loops of Henle (LOH), Descending thin limb of LOH (DTL-LOH), distal tubular cells (DT), and nephron progenitor cells (NPC), vascular cells (endothelial cell and pericyte), immune cells (macrophage, B cell and NKT cells) and proliferating cells were detected (Fig.7D, a). Marker genes were identified according to the CellMarker2.0 dataset (http://www.bio-bigdata.center/index.html). Marker genes included *FABP4, VAT1, CD68* for macrophages; *COL1A1, COL3A1, COL1A2, PDGFRA* for interstitial cells; *RAMP2, FLT1, DLL4, ALDOB* for endothelial cells; *GPX3, GLYAT, PLPP3* for developing nephrons; *RGS5, ACTA2, MYH11, TAGLN* for pericytes; *UBE2C, HMGB2, UBE2S, TUBB4B* for proliferating cells; *HLA-DQB2, HLA-DQA1, RAB30, HLA- DRB1* for B cells; *SLC12A1, UMDO, CLDN16*, *CLDN10* for LOH; *CRYAB, FST, LYPD2* for DTL-LOH; *LHX1, SLC12A3, GATA3* for DT; *RUNX2, RASGRP2, CYFIP2, RABGAP1L* for NKT cells; and *HSPA1A, CRABP2, ITGA8* for NPCs (Fig. 7D, b). Specific markers of each cell type were in Fig. 7D, b. ScRNA-seq analysis of implant of DKS+hPASCs+KIO showed that hPASCs vascularized bioengineered kidney have regenerated human kidney parenchymal cells, vascular cells and immune cells in rat partial nephrectomy model.

Next, to analyze the origins of human kidney parenchymal cells, vascular cells, and immune cells within the implants, correlation analyses were conducted between the scRNA-seq data from the implants and those from hPASCs or KIO.IntegrateLayers function is used to integrate scRNA-seq data from implants using human genome and hPASCs or KIO for correlation analysis, aiming to trace the cellular origins of implants. The MapQuery function predicts the original cell types (endothelial cells and immune cells) of implants by using hPASCs data as a reference dataset, revealing that the endothelial cells and immune cells (B cells, T cells, NK cells, macrophages) of implants were mainly composed of cell populations of angiogenesis 1 cluster derived from hPASCs (Fig.7E, a). As well as the renal parenchymal cells (interstitial cells, developing nephrons, LOH, DTL-LOH, DT, and NPC) were mainly from KIO populations such as DTC, podocyte, CT, IPC and interstitial cells (Fig.7E, b). Thus, vascular cells and immune cells of implants were mainly from hPASCs, but renal parenchymal cells were mainly from KIO.

Finally, scRNA-seq analysis of the DKS+hPASCs+KIO implant using the rat genome identified diverse rat cell types, including: endothelial cells 1 (*Tspan7, Clic5, Slc29a1, Tmem204*), endothelial cells 2 (*Cdh5, Sox7, Adgrl4, Cldn5*), and smooth muscle cells (*Myl9, Ppp1r14a, Acta2, Mcam*); Macrophage subsets included macrophage 1 (*Mrc1, Pla2g2d, Slc10a6, Alox5*) and macrophage 2 (*Ch25h, Slco2b1, Ltc4s, Mmp12*); An inflammatory macrophage subset expressed *RGD1305807, Fam20c, Fabp5, and Spp1*. Classical monocytes showed high expression of *Fcnb, Clcf1, Ari5c,* and *Clec4a2*. Mast cells expressed *Fcer1a, Mcpt8l*2, and *Ms4a2, Mcpt8* while plasmacytoid dendritic cells (*Cyb561a3, LOC24906, Slc7a1, Siglech*) and conventional dendritic cells (*Slamf7, Ctse, Clec9a, Cd226*) were also present. T cells expressed *Ncaph, Pttg1, Psmc4*, and *Stmn1*; NK cells expressed *Nkg7, Klrd1, Klre1*, and *Ccl5*; neutrophils exhibited high levels of *S100a9, S100a8, Pglyrp1,* and *Mmp8* (Extended Data Fig. 7A, B). Rat marker genes were identified according to the report before^48^. In the implant of bioengineered kidney, rat cells were predominantly immune and endothelial, but no kidney parenchymal cells.

In conclusion, the implantation of bioengineered kidneys facilitated the regeneration of human kidney parenchymal cells, vascular cells, and immune cells within the host. These human cells originated from hPASCs and KIO.

### hPASCs vascularized bioengineered kidney promote angiogenesis enhancing AKT pathway

RNA sequencing and gene ontology (GO) enrichment analysis revealed that pathway was up-regulated in blood vessel development in DKS+hPASCs+KIO group compared with DKS+hUVECS+KIO group, and regulation of angiogenesis in DKS+hUCMSCs+KIO (Extended Data Fig.7A, a). Kyoto Encyclopaedia of Genes and Genomes (KEGG) analysis further suggested that there was upregulation of the PIP3/AKT signalling pathway in DKS+hPASCs+KIO group compared with DKS+hUVECS+KIO group (Extended Data Fig.7A, b), indicating that hPASCs activated the angiogenesis related pathway after transplantation. Wester blotting further validated that in the renal subcapsular transplantation model, the expression of pAKT was upregulated in DKS+hPASCs+KIO group than DKS+hUVECs+KIO (*n*=3, *P*=0.0246) and DKS+ hUCMSCs+KIO (*n*=3, *P*=0.0032); the expression of VEGFA was upregulated in DKS+hPASCs+KIO group than DKS+hUVECs+KIO group (*n*=3, *P*=0.0398) (Extended Data Fig.7B, a). In the partial nephrectomy and bioengineered kidney-sutured rat model, the expression of pAKT was upregulated more in DKS+hPASCs+KIO group than DKS+hUVECs+KIO group (*n*=3, *P*=0.0297); the expression of VEGFA was upregulated in DKS+hPASCs+KIO group than DKS+hUVECs+KIO (*n*=3, *P*=0.0456) and DKS+hUCMSCs+KIO group (*n*=3, *P*=0.0044) (Extended Data Fig.7B, b). Accordingly, hPASCs promoted angiogenesis by activating the AKT signal pathway.

## Discussion

The present study demonstrated that hPASCs induced more vasculargenesis than that of hUCMSCs and hUVECs in the host. Firstly, hPASCs exhibit angiogenic potential: scRNA-seq revealed enriched angiogenic signatures across clusters. In 3D “Sandwich” *in vitro* models, hPASCs autonomously formed structured vascular networks, unlike hUCMSCs or hUVECs, which showed spheroidal aggregation. Co-cultures of hPASC and hUVEC recapitulated network formation, underscoring the essential role of hPASCs in vascularization. Secondly, shear stress from circumfusion promotes hPASC differentiation into hECs, facilitating vascular network formation in DKS. To mimic *in vivo* conditions, hPASCs were pre-seeded and cultured under circumfusion for 4 days, simulating arterial-to-venous flow. Optimized flow rates improved cell distribution, reduced lumen clogging, and prevented scaffold damage. Shear stress also induced endothelial differentiation and enhanced function^49^, leading to effective adhesion and expression of CD31 and α-SMA. Thirdly, hPASCs exhibited strong adhesion to the DKS, which is vital for cell survival post-transplantation. Conversely, hUCMSCs tended to aggregate and showed limited adhesion, likely due to higher traction forces. Although hUVECs adhered to the DKS, abnormal CD31 localization indicated impaired endothelial function. hUVECs weak adhesion resulted in detachment under blood flow, leading to vessel occlusion^21^ and apoptosis^50^, particularly on poorly laminin substrates^51^. Disruption of cell-matrix adhesion of ECs causes loss of anchorage and anoikis^26, 52^. Fourthly, the serious inflammatory cell infiltration in the implant is due to the strong immunogenicity of hUVECs and the activation of host immune rejection. Allogeneic antigens of ECs are a preferred target of antibodies following allogeneic transplantation^53^. Allogeneic human ECs express MHC class I and II molecules and also constitutively express costimulatory factors such as PD-L1, enhancing the direct allogeneic reaction between the ECs and T cells, ultimately leading to strong immune rejection^20^. Overall, hPASCs are the more suitable cell source for revascularizing the DKS than that of hUCMSCs and hUVECs.

*in vivo* transplantation showed the glomerular-like structures in DKS+hPASCs+KIO group contained intact basement membrane and bowman’s capsule, highly resembling native glomeruli. Firstly, we speculated that seeding KIO into hPASCs vascularized DKS *in vitro* would benefit the survival and vascularization of KIO and connect to host circulation system *in vivo*, fluorescent tracking also proved the involvement of hPASCs in building glomerular capillaries. The survival of bioengineered kidneys after transplantation entails sufficient oxygen and nutrient supply based on timely connection to host circulation system; Avascular implant is deficient in blood, oxygen and nutrient at early stage but slow extension of host vascular network into the implant will result in impaired implant integrity^54^. The maturity of KIO obtained by *in vitro* culture is similar to that of the embryonic kidney, the absence of entrance and exit of the vascular system limits the in vitro reproduction of glomerular filtration function; Despite a small number of endogenous ECs in KIO^34^, their limited quality are unable to construct a vascular system that supports the survival and function of the whole organoid, and *in vivo* vascularization after transplantation mainly relies on host-derived cells^55^. Secondly, DKS provides the natural structure, molecular microenvironment and renal-specific ECM to improve KIO maturity. Organotypic higher-order structure with collecting tubes connected to nephrons and parenchyma surrounded by stromal cells is obtained when nephron progenitors and ureteral buds are integrated with pluripotent stem cell derived lineage-specific stroma^56^.*In vitro* culture of hiPSC-derived KIO with dECM obtains highly vascularized KIO with mature glomerular development patterns by stimulating the endogenous ECs of hiPSC-derived KIO to infiltrate into glomerular-like structures^57^. Lastly, KIO contains various components such as renal stem cells, glomerulus, endothelial cells, collecting tubular, distal tubular, and proximal tubular cells; which were contributed to regenerate the kidney. In conclusion, hPASCs supported the survival and development of KIO.

Kidneys of large mammals such as pigs are more similar to human kidneys in physiology, organ scale, and embryonic development patterns than rodents, resulting in a higher clinical application value^58^. Notably, one study reported that a genetically modified pig kidney transplanted into a brain-dead patient with acute kidney injury and stage 2 chronic kidney disease-maintained creatinine clearance function and normal histological morphology^59^. However, there were still potential risks, such as xenogeneic acute rejection and animal-derived viruses affecting the human recipient^60^.This study was based on the xenotransplantation of human vascular-forming cells and human-derived kidney organoids in recipient rats. Therefore, the immunomodulatory effects of hPASCs may reduce immune rejection. In the future, we expect to conduct further research using pig models. The increased demand for cell numbers and changes in recipient species will bring about new challenges.

## Limitation of the study

In this study, an orthotopic renal transplantation model was employed, utilizing a in rat partial nephrectomy model. Due to the compensatory function of the remaining kidney, the evaluation of the filtration and reabsorption functions of the hPASCs vascularized bioengineered kidneys was limited. Future research will consider performing a complete nephrectomy to more accurately assess the urine production capacity of the bioengineered kidneys.

## Materials and Methods

### Animals for research

SRG (Sprague Dawley-*Rag2^em2hera^Il2rg^em1hera^/HblCrl*) rat (9weeks, 200g weight) and Male Sprague Dawley (SD) rat (8– 10 weeks, 250–300g weight) were purchased from Zhejiang Vital River Laboratory Animal Technology Co., Ltd., were housed at constant temperature (24 ± 1°C), and humidity (50–60%), in a specific pathogen-free grade environment with free access to food and water. All animal experiments were approved by the Ethics Committee of Wenzhou Medical University (ethical code: wydw2023-0396).

### Culture of hPASCs

The placenta was collected in a sterile bag and transported to the laboratory at 4°C. Upon arrival, it was washed thoroughly with PBS containing 2% Pen Strep to remove blood residues. Sterilized scissors and serrated forceps, pre-treated via high-pressure sterilization, were used to section the placental tissue. The segments were enzymatically digested with 1 mg/mL collagenase II (Gibico, 171701015) and IV (Gibico, 17104019) at 37°C, with gentle inversion every 10 minutes. Digestion was halted by adding 10% FBS-supplemented D/F12 culture medium, and the process was stopped early if significant tissue digestion occurred before 60 minutes. Undigested fragments were removed by sedimentation, and the supernatant was filtered through 70µm and 40 µm cell strainer (BD) to obtain microvascular tissue of diameter between 40 µm and 70µm. The suspension was centrifuged at 800g for 10 minutes, and the pellet was resuspended in pericyte medium (ScienCell^TM^, Cat.1201). Cells were seeded in sterile Petri dishes (Jet Biofil, TCD000100) and incubated in a 5% CO2 incubator at 37.5°C, with medium renewed every two days and passaging every 4–5 days. Human umbilical vein endothelial cells (hUVECs) were isolated and cultured in Endothelial Cell Medium (ScienCell^TM^, Cat.1201), while human umbilical cord mesenchymal stem cells (hUCMSCs) were isolated from Wharton’s Jelly and cultured in MSC medium (ScienCell^TM^, Cat.7501). All the cells were used within the first three passages to maintain optimal phenotypic and functional characteristics, with all experiments conducted by the third passage. The collection and use of human placenta and umbilical cord were approved by the Ethics Committee of Xuanwu Hospital of Capital Medical University (Ethical Approval Number: Linyanshen [2021]147, Approval date: 8/9/2021).

### Vascular capacity of hPASCs on 3D and 3D “Sandwich” culture

#### 3D Culture Procedure

Matrigel (R&D Systems; 3433-005-01) was thawed on ice at 4°C. A 48-well plate was preheated to 37.5°C, and 1.5 mL micro-centrifuge tubes and P200 pipette tips were precooled to 4°C. The 3D culture system was structured in two layers. The bottom layer comprised a mixture of 50% Matrigel (V/V) and 50% endothelial cell (EC) culture medium (V/V), which was incubated in a 5% CO2 incubator at 37.5°C for 30 minutes to allow solidification. The upper layer consisted of cells (5.3×10^6^ cells/mL) suspended in EC culture medium.

#### 3D “Sandwich” Culture Procedure

Fifty microliters of EC Medium and 50 microliters of Matrigel were mixed in a pre-cooled 1.5 mL microcentrifuge tube. The mixture was then transferred to a low-attachment, pre-warmed 48-well plate and incubated in a 5% CO₂ incubator at 37.5°C for 30 minutes to allow the Matrigel to solidify. Six types of cell suspensions were prepared: hUVECs, hUCMSCs, hPASCs, hUVECs+hUCMSCs, hUVECs+hPASCs, and hUVECs+hUCMSCs+hPASCs. Next, 75 microliters of each cell suspension were mixed with 75 microliters of Matrigel (1:1, V/V) and transferred to the 48-well plate. This mixture was incubated in a 5% CO₂ incubator at 37.5°C for another 30 minutes to solidify the Matrigel. Finally, 250 microliters of pre-warmed Endothelial Cell Medium were added, and the culture medium was replaced daily.

### Live-cell imaging

hPASCs at passage 3 were cultured in 3D Matrigel and imaged using a biostation CT (Nikon) at 37°C with 5% CO2. Cell images were captured every 2 hours and recorded continuously for 96 hours. The images were then exported to a computer for analysis and compilation into dynamic videos.

### Differentiation ECs and SMCs of hPASCs

hPASCs and hUCMSCs were seeded at a density of 3, 000 cells/cm² in cell culture flasks and cultured in EGM-2 medium (Lonza, CC-3156) supplemented with 10 μM SB431542. The medium was refreshed every two days, and the differentiating endothelial cells (ECs) were passaged every 4–6 days. The culture was maintained for a total of 14 days. For smooth muscle cell (SMC) differentiation, both hPASCs and hUCMSCs were cultured in DMEM/F12 medium containing 10% fetal bovine serum (FBS) (Gibico, A5669701) for three passages.

### Flow cytometry identification of hPASCs

Antigen-labelled hPASCs were analysed using flow cytometer (BD, Canto II). Cellular information was analysed using Flow Jo-V10 (14.0.0.0). Antibodies used in flow cytometry were listed as following. Mouse Anti-Human CD201 APC (EPCR) (eBioscience™, 17-2018-42), Mouse Anti-Human CD276 APC (eBioscience™, 17-2769-42); Human/Mouse CD44 PE-Cyanine7 (eBioscience™, 25-0441-81); Mouse Anti-Human CD34 PE (BD Pharmingen™, 550761), Mouse Anti-Human CD146 PerCP-Cy™5.5 (BD Pharmingen™, 562134), Mouse anti Human CD31 PE (BioLegend, 102418), Mouse Anti-Human CD326 APC (BioLegend, 118214).

### Traction force assay of hPASCs

The glass bottom of a 35 mm confocal dish was pretreated with 3-aminopropyltriethoxysilane (APTES, Sigma) for 15 minutes, followed by 0.5% glutaraldehyde in PBS for 30 minutes to promote polyacrylamide (PA) hydrogel attachment. A mixture of acrylamide and bis-acrylamide was prepared at a ratio of 5% to 0.1%, incorporating 1% fluorescent beads (0.2 μm diameter, Thermo Fisher, F8810) to create PA gels with a stiffness of 3 kPa. The gel was activated with Sulfo-SANPAH (Pierce) and coated with 0.2 mg/mL collagen I at 4°C overnight. Subsequently, cells were seeded onto the substrate overnight to ensure proper adhesion. Phase contrast images of single cells and fluorescent images of the beads were captured before and after cell digestion. Digital image correlation (DIC) was employed to calculate displacement fields of the substrates resulting from cell contraction. The cell traction stress field was reconstructed using the optimal filtering method based on the classic Fourier Transform Traction Cytometry (FTTC) technique, implemented in MATLAB.

### 3D “sandwich” culture of human fetal kidney organoids

The primary cells used in this study were derived from human fetal kidneys (16–18 weeks). Tissue was treated with a 1:1 mixture of collagenase I (Gibco, 171018029) and collagenase II (Gibco, 171701015) at a concentration of 1 mg/mL. Digestion was conducted at 37°C for 30 minutes and was terminated by adding DMEM/F-12 (Sigma; D8537-500 mL) supplemented with 10% fetal bovine serum (FBS). The cells were then centrifuged at 800 × g for 5 minutes to pellet the cell fractions. The study involving donated human fetal tissue samples received ethical approval from the Ethics Committee of the Eye Hospital of Wenzhou Medical University (Project: A Comprehensive Study on the Generation and Utilization of Fetal-Derived Organoids; Research License: 2025-137-K-109-01).

The human kidney organoids culture medium (hKOCM) comprised pericyte medium(termed PM-hKOCM, including 1% Pen Strep (V/V), 1% GlutaMAX-1 (V/V), 1% MEM NEAA (V/V) (Gibco, 11140-050), 10 mM HEPES (Solarbio, H1095), 1% Insulin-Transferrin-Selenium-X (V/V) (Gibco, 51500-056), 0.1% 2-mercaptoethanol (V/V) (Gibco, 21985-023), 2% B-27 (V/V) (Gibco, 17504044), 1% KnockOut SR XenoFree medium (KSR) (Gibco, 12618013), 100 ng/mL R-spondin 1 (R&D systems, 4645-RS-025), 25 ng/mL Wnt 3a (R&D systems, 5036-WNP), 3 µM CHIR-99021 (Selleck, S1263), 50 ng/mL EGF (Peprotech, 100-47), 50 ng/mL FGF10 (Peprotech, 100-26), 50 ng/mL KGF (Peprotech, 100-19), 0.2 µM A 83-01 (MedChemExpress, HY-10432) 50 ng/mL GDNF (Peprotech, 450-10), 0.2 µM LDN193189 (MedChemExpress, HY-12071), 5 µM SB 202190 (MedChemExpress, HY-10295), 0.1 µM TTNPB (MedChemExpress, HY-15682), 10 µM Y-27632 (Selleck, S6390), 0.2% (V/V) Primocin™ (V/V) (Invivogen, ant-pm). Primocin™ and Y- 27632 are added only during resuscitation and passage. Another medium using Advanced DMEM/F-12 (Gibco, 12634- 010) with same supplement to PM-hKOCM(termed D/F12-hKOCM.

Before the 3D “sandwich” culture, the 48-well plate (Jet Biofil) was preheated in a CO₂ incubator. The PM-hKOCM was pre-chilled and supplemented with 10 µM Y-27632 (Selleck; S6390) and 0.2% Primocin™ (V/V) (Invivogen, ant-pm). Matrigel (R&D Systems; 3433-005-01) was kept on ice to preserve its integrity.The 3D “sandwich” culture system consisted of three layers. The bottom layer was a mixture of 50% Matrigel (V/V) (R&D Systems, 3433-005-01) and 50% PM-hKOCM (V/V), which was added to the preheated plate and gently tilted to spread the Matrigel. This layer was then solidified at 37°C in a CO₂ incubator for 30 minutes. The middle layer was composed of 30% Matrigel and 70% PM- hKOCM, into which the primary kidney cells were mixed. After thorough mixing, this layer was added on top of the solidified bottom layer and allowed to solidify at 37°C for 30 minutes. Finally, 300 µL of PM-hKOCM was slowly added on top of the solidified middle layer. The entire construct was placed in a CO₂ incubator, with daily medium changes.

### Passage of human kidney organoid

After 5–6 d of primary renal organoid culture, 200 μl of pre-chilled DMEM/F12 was introduced into each well of the culture plate. The organoids, along with the gel mixture, were initially transferred to a 1.5 mL EP tube using a 1 mL pipette tip with a slanted cut. The organoids were then repeatedly pipetted up and down to promote dispersion. Subsequently, the mixture was pipetted 2–3 times using a 1 mL sterile syringe (KDL) and centrifuged at 500 ×*g* for 5 min to separate the organoids from the Matrigel. The supernatant was carefully aspirated multiple times using a 200 μl pipette tip, and then 200 μl of 0.25% Trypsin-EDTA (Gibco; 25100-072) was added to the tube. Digestion was allowed to proceed at 37°C for 3 min. Next, the mixture was pipetted 2–3 times using a 1 mL sterile syringe (KDL) and 2–3 times using a 1 mL insulin syringe to dissociate the organoids into small cell clusters. The digestion process was stopped by adding 10% FBS DMEM/F-12, after which centrifugation was performed at 350 ×*g* for 5 min. Passage of the organoids was then performed using the 3D “sandwich” culture method with daily medium changes. For long-term storage, the organoids were combined with a cryopreservation solution consisting of 60% FBS, 10% DMSO, and 40% DMEM/F12. The collected organoids were mixed with the cryopreservation solution and stored at -80°C overnight before being transferred to liquid nitrogen for long-term preservation.

### Decellularized of rat kidney scaffolds

Rats were secured in the supine position after inhalation anaesthesia with 2% isoflurane (RWD, R510-22-10). A vertical abdominal incision was made from the symphysis pubis to the xiphoid region to expose the peritoneal cavity. The kidney was flushed with 100 mL heparin solution injected through the hepatic portal vein to remove residual blood, after which the perirenal fat and renal capsules were removed. Blunt isolation of the abdominal aorta, inferior vena cava, and kidneys was performed bilaterally. The renal artery and the 1 cm abdominal aorta connected to it were preserved; the renal vein and the 1 cm inferior vena cava connected to it were also preserved. The renal artery and vein were cannulated with 24G indwelling needles, and extravascular ligation was performed using a 4-0 suture (Jinhuan Medical Products Co., Ltd.). The kidney was placed in a 10 cm sterile petri dish (JET, 02023662-TCD010100) with a beaker underneath to hold the waste liquor. Then, a peristaltic pump (Longer pump, BT300-2J) assembled with pump head (DG-4, 10 rollers) was used to perfuse the kidney with decellularization solution at a speed of 5 rpm (5 mL/min) in order: 500 mL of 1 g/L heparin solution, 500 mL of 1% (v/v) Triton X-100 solution (Beyotime, ST795), and 800 mL of 8 kg/L SDS solution. The kidney was then perfused with 2000 mL phosphate-buffered saline (PBS) mixed with 1% penicillin-streptomycin solution (Gibco, 15140-122) at 1.5 rpm (1.5 mL/min) for 1300 min and kept in a 50 mL centrifuge tube containing this solution at -80° C to remove residual perfusate.

### Recellularization of rat kidney scaffolds and whole organ culture

A total of 2×10^7^ cells were trypsinzed, centrifuged (1800 rpm, 5 min) and resuspended in a 150 μL medium containing 10% MATRIGEL. The cell suspension was equally infused into the DKS through cannulas attached to the renal artery and vein using a 1 mL syringe. The recellularized kidney scaffold was kept in a 50 mL centrifuge containing a 20 mL medium tube in a mobile phase bottle for culture. During cell culture, the arterial and venous cannulas were connected to a peristaltic pump tube, through which the medium was recirculated from the arterial side to the venous side at a speed of 17 rpm (1 mL/min). The medium was replaced on days 3 and 5, and the kidney scaffold was removed from the cell culture bioreactor.

A total of 1×10^6^ KIOs were resuspended in 150 μL medium, after which the kidney organoids were seeded and cultured as described above for 48 h. The kidney scaffold was removed from the cell culture bioreactor on day 2 for transplantation and sample collection. Samples were divided into three sections for use in paraffinization, electron microscopy, and transplantation.

### Scanning Electron Microscopy (SEM)

Samples measuring 4 mm × 3 mm × 3 mm were fixed in 2.5% glutaraldehyde (EMS, 16020) at 4°C for 12 hours. Following fixation, the samples were rinsed with phosphate-buffered saline (PBS) and stored at -80°C for 24 hours. Subsequently, the samples were dehydrated using a vacuum freeze dryer (Labconco, KS, USA) for 24 hours. After drying, the samples were coated with gold using a vacuum sputter coater (Hitachi, Tokyo, Japan) for 2 minutes. Scanning electron microscopy images were then acquired using a scanning electron microscope (Hitachi, Tokyo, Japan).

### Transmission Electron Microscopy (TEM)

Samples measuring 1 mm³ were fixed in 2.5% glutaraldehyde overnight at 4°C. Following fixation, the samples were rinsed with 0.1 M phosphate buffer (Shanghai Yubo Biotechnology Co., Ltd., YB160232) and subsequently exposed to 1% osmium tetroxide (Ted Pella, 18459) for 1 hour. The samples were then washed with phosphate buffer and distilled water. After fixation, the samples were stained with 1% uranyl acetate (Syntechem, 541-09-3) for 2 hours and dehydrated through a series of graded acetone solutions. The samples were immersed in a 1:1 mixture of acetone and epoxy resin (SPI, 1220324) and maintained at 37°C for 1 hour before being fully embedded in epoxy resin. An ultramicrotome was employed to obtain sections with a thickness of less than 100 nm. Images were captured at magnifications of 15, 000× and 30, 000× using a transmission electron microscope (Hitachi, Tokyo, Japan).

### Glomerular filtration

To evaluate the functionality of hPASCs vascularized bioengineered kidney *in vitro*, we used the method that have been reported before^15^, in which perfusion fluid was filtered and reabsorbed through cadaveric kidney, DKS, and DKS+KIO+hPASCs group. The perfusion fluid, consisting of NaCl, KCl, KH2PO4, MgSO4, CaCl2, NaHCO3, and glucose in Krebs-Henseleit buffer (Pricella, PB180348), was supplemented with bovine serum albumin (BSA) (Beyotime, ST025- 20g), creatinine (Sigma, C4255-10G), glycine (MCE, HY-Y0966), L-proline (MCE, HY-Y0252), and L-serine (MCE, HY-N0650). After filtration through a 0.22 μm mesh, the perfusion fluid was incubated in a 37°C, CO2 incubator. Perfusion was carried out via arterial cannulation at 10 rpm, and the resulting urine and venous outflow were collected and stored at -80°C. The time taken for each group to produce 100 μL of urine was recorded, and the average urine flow rate (mL/min) was calculated and compared between the cadaveric kidneys, DKS group, and DKS+hPASCs+KIO group. The urine samples were analyzed by an automated biochemical analyser (HITACHI 3500) to assess levels of glucose, creatinine, and albumin. Creatinine clearance rate (CCr), calculated as urine creatinine (mg/dL) multiplied by urine volume per minute (mL/min) divided by venous creatinine (mg/dL), was utilized to evaluate glomerular filtration rate. The fraction of solute excretion, including albumin, glucose, sodium, potassium, chloride, and calcium in urine, was employed to assess renal tubular reabsorption and secretion function.

### Renal subcapsular and partial nephrectomy transplantation

#### Renal subcapsular transplantation

After inhalation anaesthesia with 2% isoflurane, rats were secured in the right lateral decubitus position. A 1.5–2 cm vertical incision was made in the left renal region to expose both kidneys under sterile conditions. Two mm incisions were made in the inferior pole of the renal capsule, after which the grafts were implanted into the subcapsular area and pushed to the superior pole using microforces. After the kidney was put back to the abdomen, the incision was sutured with 4-0 stitches in layers. The rats were sacrificed on day 14 for sampling.

#### Partial nephrectomy transplantation

The rats were maintained in the right lateral decubitus position after being anesthetized with 2% isoflurane. The left kidney was aseptically exposed through a dorsal transverse incision (1.5–2 cm) in the renal region, whereas the right kidney was left untreated. The left renal artery and vein were isolated by blunt dissection and occluded using a nontraumatic microvascular clamp. The left kidney turned purple after clamping, and the clamping was maintained for 35 min to prevent ischemia-reperfusion injury. One-third of the inferior pole of the kidney was resected. The remaining kidney was then isometrically sutured with trimmed scaffolds of similar volume as the resected kidney using absorbable 5-0 sutures (Jinhuan Medical Products Co., Ltd., R516) at approximately 2 mm intervals with eight to nine stitches. The vascular clamps were removed to restore reperfusion after transplantation, and the kidneys immediately reverted to red. After the left kidney was returned to the abdomen, the incision was sutured in layers using nonabsorbable 4-0 sutures (Jinhuan Medical Products Co., Ltd., F402). The rats were euthanized after 14 days to harvest their left kidneys.

### Histochemistry staining

After being fixed with 4% paraformaldehyde for 24 hours, and embedded in paraffin, kidney samples were cut into 4 µm thick sections. The sections were heated at 60°C for 1 h, deparaffinized using xylene and dehydrated using a graded series of alcohol solutions and morphological characterization was performed using histochemical analysis. Haematoxylin and eosin (Solarbio, G1120) and Masson’s trichrome (Solarbio, G1343) periodic acid-Schiff (Solarbio, G1281) staining were performed according to manufacturer’s protocols.

### Immunofluorescence staining

Immerse tissue sections in PBS (Solarbio, P1010) for 10 minutes. Permeabilize with 0.25% Triton X-100 (Beyotime, P0096) for 5-15 minutes, followed by washing with PBS containing 0.1% Tween-20 (Solarbio, T8220, PBST). Block with 5% donkey serum (Solarbio, SL050) for 30-60 minutes. Incubate with the primary antibody diluted in 2.5% donkey serum overnight at 4°C. Wash with PBST, then incubate with the secondary antibody diluted in 2.5% donkey serum for 1 hour and 15 minutes. Wash with PBST, add DAPI (Solarbio, S2110), mount the slides, and examine using a laser confocal microscope (Zeiss, LSM 900). Analyse the data with ZEN 2.6.

Primary Antibodies: CD31 (Abcam; ab281583, diluted 1:100);α-SMA (Abcam; ab7817, diluted 1:150); human podocalyxin antibody (PODXL) (R&D Systems; AF1658, diluted 1:250); Lotus tetragonolobus lectin (LTL) fluorescein (Vector Laboratories; FL-1321-2, diluted 1:100-1:400);Purified mouse anti-E-cadherin (CDH1) (BD Biosciences; 610182, diluted 1:50); Rabbit anti-pax2 antibody (Abcam; ab79389, diluted 1:50);Anti-Lrp2 (Abcam; ab76969, diluted 1:200);NKCC2 polyclonal antibody (Proteintech; 18970-1-AP, diluted 1:100-1:200); Aquaporin 2 (AQP2) (Santa Cruz; sc-515770, diluted 1:250); NPHS1 Polyclonal antibody (R & D system; AF4269-SP, diluted 1:400-1:500); NPHS2 Polyclonal antibody (Proteintech; 20384-1-AP, diluted 1:400-1:500); WT1 (Santa Cruz; sc-7385, diluted 1:400); EHD Polyclonal antibody (Proteintech; 25320-1-AP, diluted 1:200); CNN1 (CST;17819T, diluted 1:500); CDH5 (Abcam; ab313632, diluted 1:500); MECOM (ATLAas antibodies; HPA046537, diluted 1:500); PAI 1 (Abcam; ab222754, diluted 1:1000).

Secondary Antibodies: Alexa Fluor®488-labeled goat anti-mouse (Abcam; ab150113, diluted 1:1500); Donkey anti-goat IgG (H&L) Alexa Fluor 488 (Abcam; ab150129, diluted 1:750); Goat Anti-rabbit IgG (H&L) FITC (Thermo Fisher Scientific; F-2765, diluted 1:750); Donkey Anti-Rabbit IgG H&L (Alexa Fluor® 594) (Abcam; ab150076, diluted 1:750); IFKine™ Red Donkey Anti-Goat IgG (Abbkine; A24431, diluted 1:150); Donkey Anti-Goat IgG H&L (Cy5) (BIOSS; bs-0294D-Cy5, diluted 1:750); Donkey Anti-Sheep IgG (H+L) Alexa Fluor™ 647 (Thermo Fisher Scientific; A21448, diluted 1:750).

### B-scan ultrasonography

A high-resolution animal ultrasonography system (VisualSonics, Vevo 3100) was used to perform kidney B-scan ultrasonography on day 14 after partial nephrectomy and transplantation. After anesthetizing rats with 2% isoflurane and placing them the right lateral decubitus position, an MX250: 20 MHz probe was used to examine the left renal region. The probe was moved slowly from the ventral side to the dorsal side near the backbone, parallel to the axis of the kidney, to scan the entire kidney. The size, shape, location, and blood flow of the host kidneys and transplantation areas were recorded.

### Single-cell RNA sequencing and Analysis

hPASCs or Cells from P3 KIO and kidney implants were disaggregated using enzymatic and mechanical methods, then filtered through a 40 μm mesh to isolate cells for single-cell analysis. A total of 10,000 cells were used to construct a single-cell nuclear transcription library with the Single-cell Sequencing Library Construction System (Singleron Matrix, Singleron Biotechnologies) and the DaSCOPE Single Cell Dynamic RNA Library Kit (Singleron Biotechnologies). Following library construction, Illumina sequencing analysis was performed.

Single-cell sequencing data were obtained from human fetal kidney organoids (in this study). CeleScope (version 2.0.4) (https://github.com/singleron-RD/CeleScope.Github) was used to process the single-cell transcriptome data and align them with the human reference genome assembly hg38^61^. Quality control screening was performed on the cells using Seurat (version 4.4.0)^62^, resulting in the removal of cells with fewer than 200 detectable genes, fewer than 500 gene counts, or a mitochondrial ratio exceeding 10%. High-quality cells were log-normalized and scaled using Seurat workflow and the top 2000 highly variable genes (HVGs) were identified using the FindVariableFeatures function in Seurat. Principal component analysis (PCA) was performed to extract 30 principal components (PCs) that explained the maximum variance among the top 2000 HVGs. Subsequently, high-quality cells were visualized in two-dimensional space using uniform manifold approximation and projection (UMAP). FindMarker function using Wilcoxon rank-sum test in Seurat was used to identify the differentially expressed genes (DEGs) in one cluster compared with other clusters. ClusterProfiler (version 4.6.2) package is used for Gene Ontology (GO) and Kyoto Encyclopedia of Genes and Genomes (KEGG) pathway enrichment analysis^63^. Kidney gene expression patterns were collected and used for cell type annotation. Monocle3 (version 1.3.1) was used to perform trajectory analyses to predict the differentiation paths between each fetal kidney organoids cell type without prior knowledge of differentiation time or direction^64^. While it is difficult to determine the starting point of differentiation without prior knowledge, package CytoTRACE (version 0.3.3) can help predict the order of cell differentiation states and stemness of cells^65^. CellphoneDB (version 3.0.0)^66^ is an open-access database that meticulously selects receptor, ligand, and their interaction information from databases such as UniProt, Ensembl, PDB, IUPHAR, etc., allowing for a comprehensive and systematic analysis of intercellular communication molecules and the study of interactions and communication between different cell types. By invoking the statistical analysis of the CellphoneDB software package, the ligand-receptor relationships characterized in single-cell expression profiles were analysed. IntegrateLayers function is used to integrate scRNA-seq data for correlation analysis; furtherly, the MapQuery function predicts the original cell types.

### RNA-sequencing analysis

Tissue samples (20–50 mg) were placed in RNase-free EP tubes (Axygen, MCT-150-C) and stored at -80°C. Samples were homogenized in 1 mL TRIzol (Ambion, 15596026) for 50 seconds at 60 Hz using a Tissue Grinder (Shanghai Jingxin, JXFSPTPR-24), incubated at room temperature for 5 minutes, and the homogenate was transferred to a new tube. Chloroform (200 μl, Aldrich, 485403-500MG) was added, mixed, incubated for 3 minutes, and centrifuged (12, 000 rpm, 15 min, 4°C). The aqueous phase was collected, supplemented with 5 μl Acryl Carrier (Solarbio, SA1020) and 500 μl isopropanol (Titan, G75885B), mixed, incubated for 10 minutes, and centrifuged (12,000 rpm, 10 min, 4°C). The supernatant was discarded, and 1 mL of 75% ethanol was added, mixed, and centrifuged (7,500 rpm, 2 min, 4°C). The remaining supernatant was dried for 5 minutes and resuspended in 30 μl RNase-free dH2O (Takara, RR047A). RNA concentration and quality were assessed using a Nano Drop One (Thermo Fisher Scientific), targeting A260/A280 ratios of 1.9-2.0. Total RNA purity and integrity were further evaluated with a Kaiao K5500 spectrophotometer and RNA Nano 6000 Assay Kit (Agilent Technologies, 5067-1511). mRNA was purified using Poly-T oligo-attached magnetic beads and fragmented with divalent cations. The mRNA fragments served as templates for first-strand cDNA synthesis, followed by complementary strand synthesis to form double-stranded cDNA. Library fragments were purified with the QiaQuick PCR Purification Kit (Qiagen, 28104), terminally repaired, A-tailed, and ligated with adapters. Target products were amplified via PCR to construct the library. The RNA library was clustered using the HiSeq PE Cluster Kit v4-cBot-HS (Illumina) and sequenced on the Illumina platform, yielding 150 bp paired-end reads. Post-sequencing, enriched differential gene expression was analyzed for highly expressed signaling pathways, western blotting used for validation. Three samples were replicated, and a *P* value of 0.05 was considered significant.

### Western blotting

Kidney tissues of implants were lysed with RIPA buffer (Beyotime, P1048), homogenized, and incubated in ice-water for 10 minutes. The supernatant was obtained via centrifugation (4°C, 12,000 rpm, 10 min). BCA working and standard solutions (ThermoScientific, 23225) were prepared, and optical density was measured at 562 nm to establish a standard curve for sample concentration calculation. A 5×loading buffer (Fude Biological Technology, FD002) was added to samples (1:4 v/v), boiled at 99°C for 10 minutes, and stored at -20°C. Samples and markers (ThermoScientific, REF26617) were loaded onto PAGE gels (YAMAY BIOTECH, PG21) and electrophoresed at 80 V for 30 minutes, then at 100 V until bromophenol blue reached the bottom. PVDF membranes (Immobilon, ISEQ00010) were activated with methanol (Hongsheng Fine Chemical, CY20211010) for 5 minutes and transferred at 300 mA for 1.5 hours. Membranes were incubated in 5% Blotting Grade (Beyotime, P0216) for 1 hour at room temperature and with primary antibodies overnight at 4°C. The primary antibodies used included AKT Rabbit pAb (abclonal, A11016), Phospho-AKT Rabbit pAb (abclonal, AP0274), VEGFA Rabbit pAb (abclonal, A5708), and β-Actin Rabbit mAb (abclonal, AC026) as an internal control. Secondary antibodies (Gour anti-rabbit IgG, HRP-linked Antibody, CST, 7074) were incubated at room temperature for 1 hour. Images were captured using a multifunctional gel imaging system (BIO-RAD, USA), and gray values were analysed with Image Lab and ImageJ 3.0.

### Statistical analysis

Statistical analyses were conducted using IBM SPSS Statistics 25, with graphs generated in GraphPad Prism (Version 8.0.1). Normality was assessed via skewness and kurtosis values; data were considered normally distributed if the absolute values were <1.96. Quantitative data with normal distribution are presented as mean ± standard deviation, with Levene’s test verifying homogeneity of variance (*P* > 0.1). Independent two-sample t-tests were applied for samples with homogeneous variance, while Welch’s t-test was used for unequal variance. One-way ANOVA with Tukey’s post-hoc test compared multiple groups with homogeneous variance, and Tamhane’s T2 test was used for those with unequal variance. Non-normally distributed data are reported as median and interquartile range, with α set at 0.05. Mann–Whitney U tests and Kruskal–Wallis tests were used for comparisons of two and multiple groups, respectively, with statistical significance defined as *P* < 0.05.

## Reporting Summary

Further information on research design is available in the Nature Research Reporting Summary linked to this article.

## Data availability

The main data supporting the results in this study are available within the paper and its supplementary information. Gene- expression data obtained by RNA-seq are available from the NCBI Gene Expression Omnibus, via the accession number. The raw and analysed datasets generated during the study are available for research purposes from leading corresponding author. Source data are provided with this paper.

## Acknowledgements

Funding was provided by the National Key Research and Development Program of China (Xuguang Du, 2023YFC3404301); Zhejiang Provincial Natural Science Foundation of China (Junjun Xu, LY24H180005); Open Project of State Key Laboratory of Animal Biotech Breeding (Junjun Xu, 2025SKLAB6-16);The Summit Advancement Disciplines of Zhejiang Province (Wenzhou Medical University–Pharmaceutics); Zhejiang Provincial Natural Science Foundation of China (Hui Liu, LZ24H120003); We also thank Scientific Research Centre of Wenzhou Medical University for consultation and instrument availability that supported this work.

## Author contributions

H.L., X.W., Y.S., Y.Z., X.R., W.S., X.Z., Y.W., J.H., Y.M., J.Z., Y.C. S.L., and F.L. performed the experiments; Y.G. contributed to other resources of the experiments; X.D. contributed to other resources of the experiments and supervised the project; J.X. conceived the hypothesis, designed and supervised the project; all authors read the manuscript and agree with its contents.

## Competing interests

Junjun Xu is inventor on two patents (application number: 02510757426X; 20251075766212) related to the content of this article and the authors declare no conflict of interest.

## Correspondence and requests for materials

All data should be addressed to the lead correspondence author Junjun Xu.

## Additional information

**Extended Data Fig.1.**
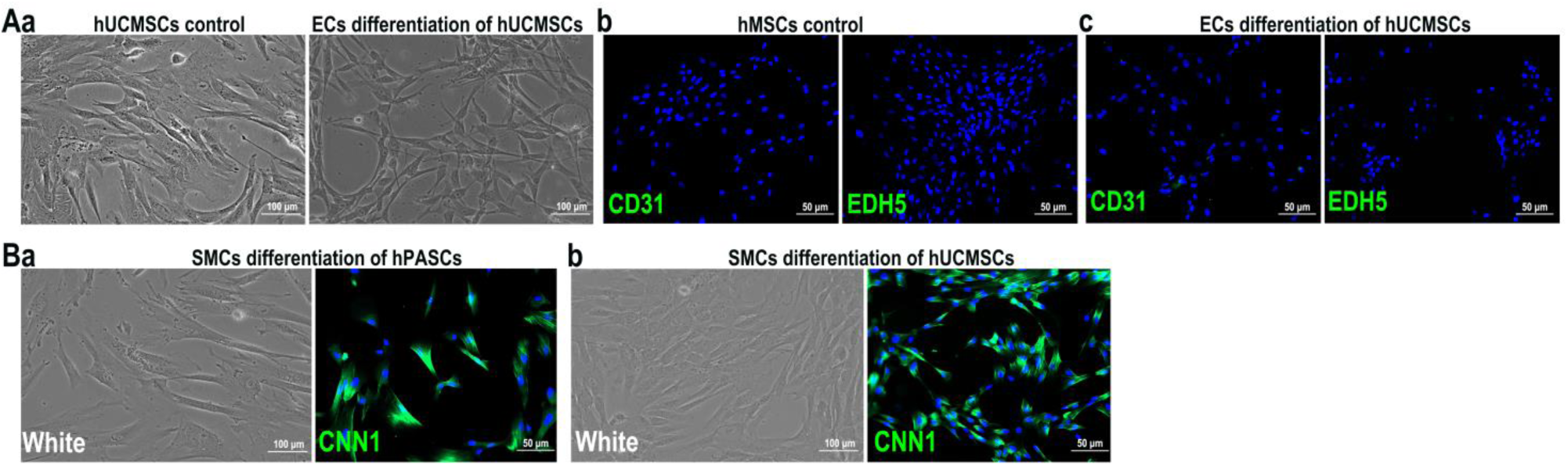
Vascular differentiation of hUCMSCs: (A), ECs differentiation: (a), Morphology of ECs differentiation of hUCMSCs; (b-c), Both hUCMSCs and post-differentiation of hUCMSCs remained negative for CD31 and CDH5. (B), SMCs differentiation: (a), SMCs differentiation of hPASCs were positive for SMC marker CNN1; (b), SMCs differentiation of hPASCs were positive for SMC marker CNN1. Scale bar 100 μm (White);50 μm (Fluorescence).

**Extended Data Fig.2.**
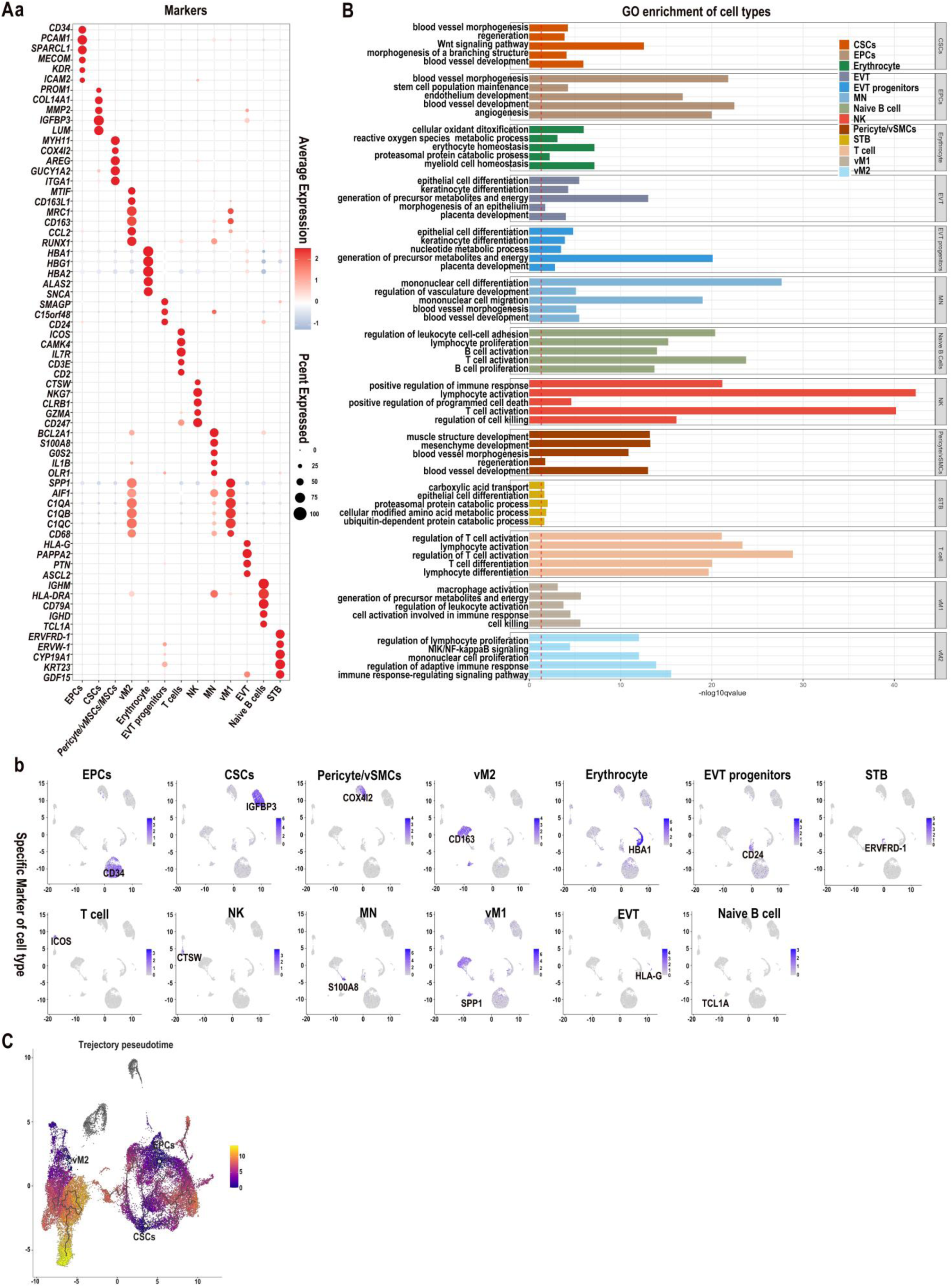
scRNA-seq analysis of microvessels: (A), ScRNA-seq analysis: (a), Marker genes of each cell type of placental microvessles; (b), Specific marker genes for each cell type. (B), GO enrichment analysis further characterized each cell type. (C), Trajectory pseudotime analysis between hPASCs and placental microvessles.

**Extended Data Fig.3.**
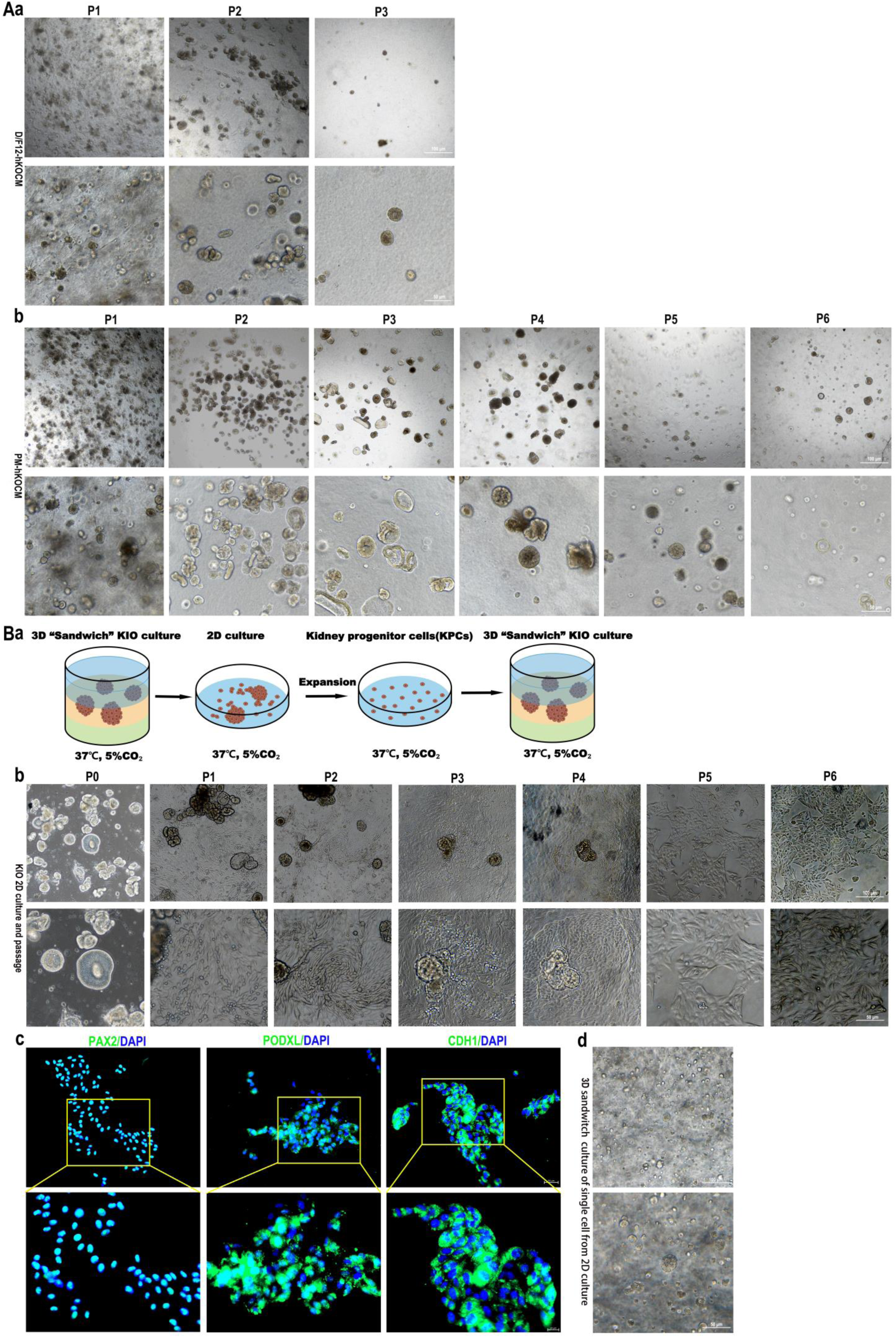
Culture and amplification of KIO of 3D “sandwich” system. (A), Ddifferent culture system of KIO: (a), The passage of KIO of culture medium with D/F12-hKOCM; (b), The passage of KIO of culture medium with PC-hKOCM. Scale bar 100 μm (up); 50 μm (down); (B), KIO from 2D culture: (a) Scheme of culture and amplification of KIO from 3D “sandwich” to 2D; (b), Multi-passage of KPCs. Scale bar 100 μm (up); 50 μm (down); (c), Iimmunofluorescence of KPCs. Scale bar 50 μm (d), Culture of KIO from 2D to 3D “sandwich”. Scale bar 100 μm (up); 50 μm (down).

**Extended Data Fig.4.**
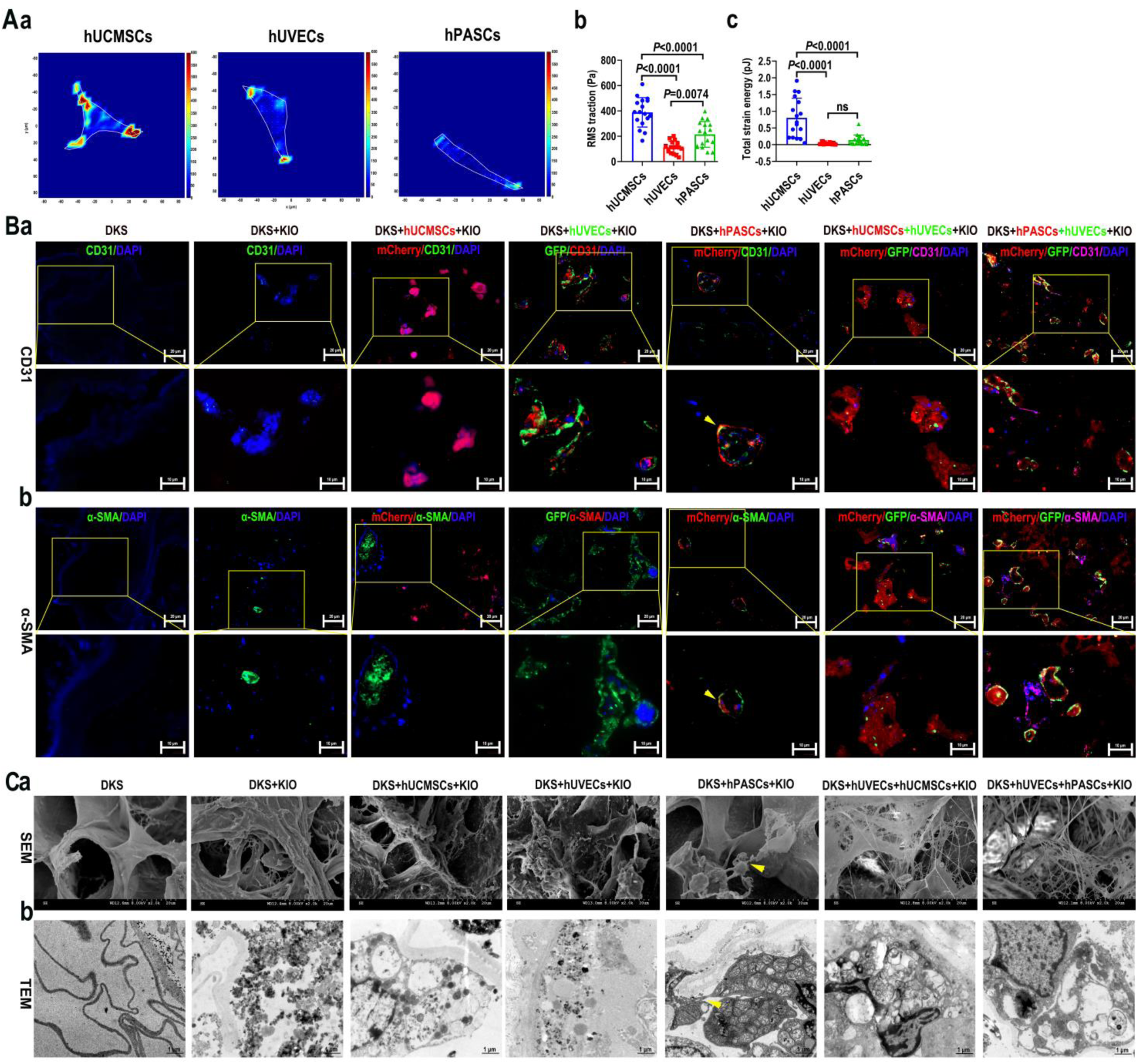
Vascularization of DKS with hPASCs *in vitro.* (A), Traction force assay: (a), Single cell TFM; (b), Single cell RMS; (c), Single cell total strain energy (pJ); *n*=17(*P*<0.05. (B), (a-b) Immunofluorescence of CD31 and α-SMA showed hPASCs reconstructed vascular structures (arrowhead). Scale bar 100 μm (up); 50 μm (down). (C), Ultrastructure of vascularized bioengineered kidney: (a), SEM showed that hPASCs formed connections between cells (arrowhead). Scale bar 20 µm; (b), TEM also showed that hPASCs formed connections (arrowhead). Scale bar 1 µm.

**Extended Data Fig. 5.**
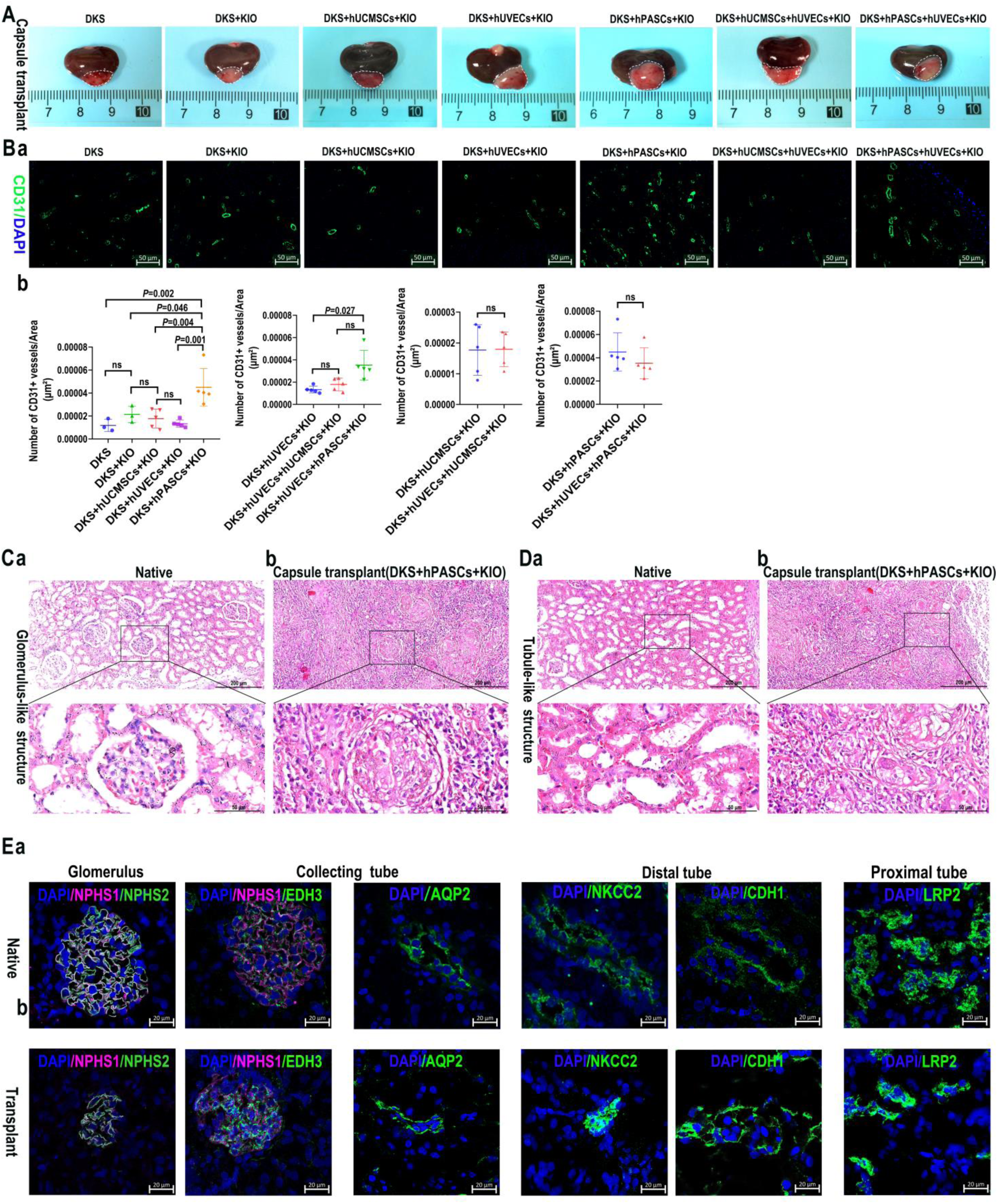
The structure and promotion vascular density of hPASCs vascularized bioengineered kidney in renal subcapsular transplantation. (A), The photograph of bioengineered kidney 14 days post transplantation; (B), hPASCs promote angiogenesis: (a), Immunofluorescence of CD31 staining; (b), Statistical analysis of the density of vascular structures; (c) The immunofluorescence co-staining results of EHD3, α-SMA, and CD31 with hPASCs. (C), HE staining analysis of Glomerulus-like structure: (a), Native group; (b), DKS+hPASCs+KIO implant group. Scale bar 200 μm (up); 50 μm (down); (D), HE staining analysis of Tubule-like structure: (a), Native group; (b), DKS+hPASCs+KIO implant group. Scale bar 200 μm (up); 50 μm (down); (E), Fluorescent staining of implant: (a), The native tissue was as control; (b), Surface markers of glomerulus (NPHS1, NPHS2), collecting tube (AQP2), distal tube (NKCC2, CDH1), and proximal tube (LRP2); Scale bar 20 µm.

**Extended Data Fig.6.**
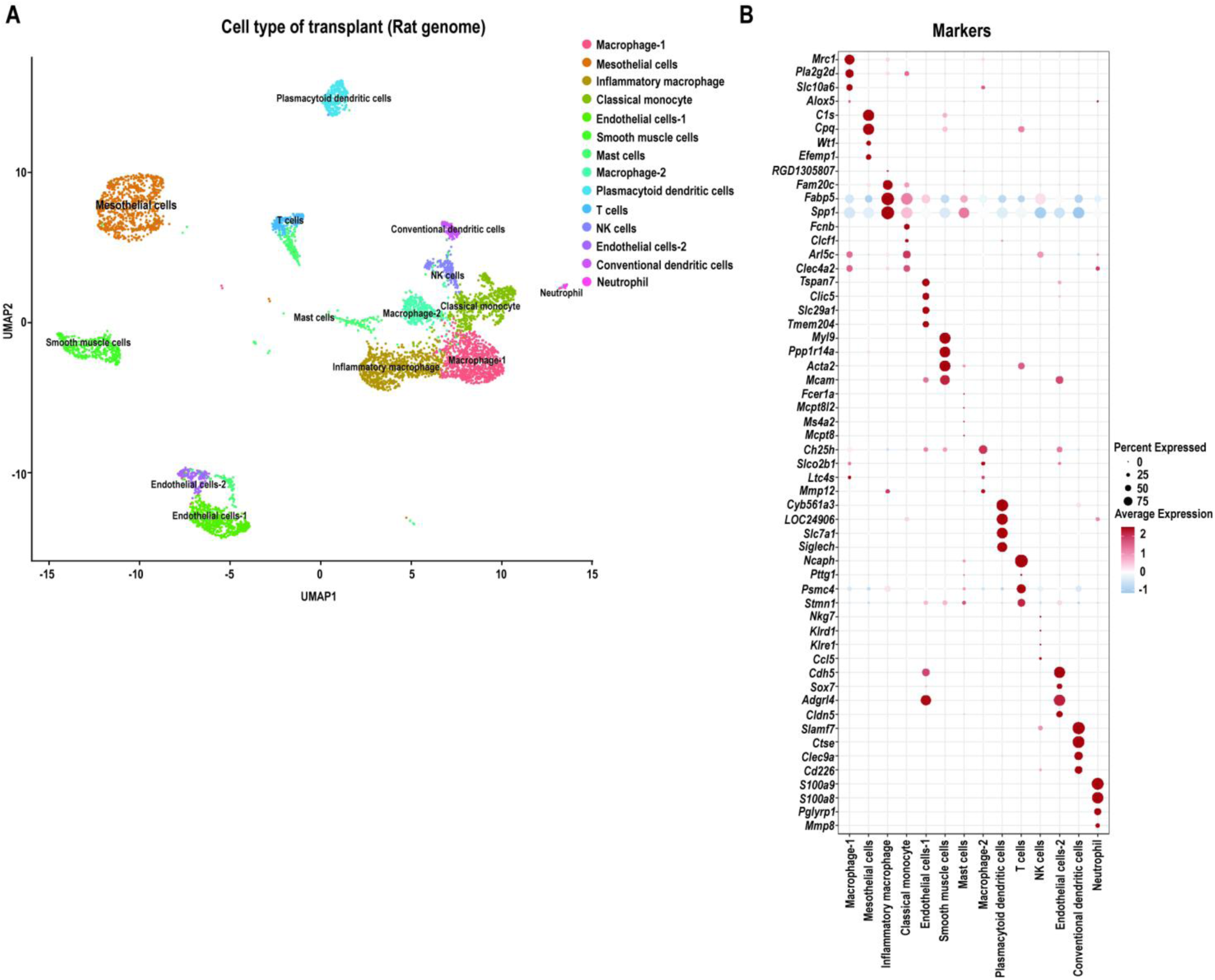
ScRNA-seq analysis of implant of DKS+hPASCs+KIO using rat genome: (A) Rat cell types of transplants; (B) Markers genes of each cell type.

**Extended Data Fig.7.**
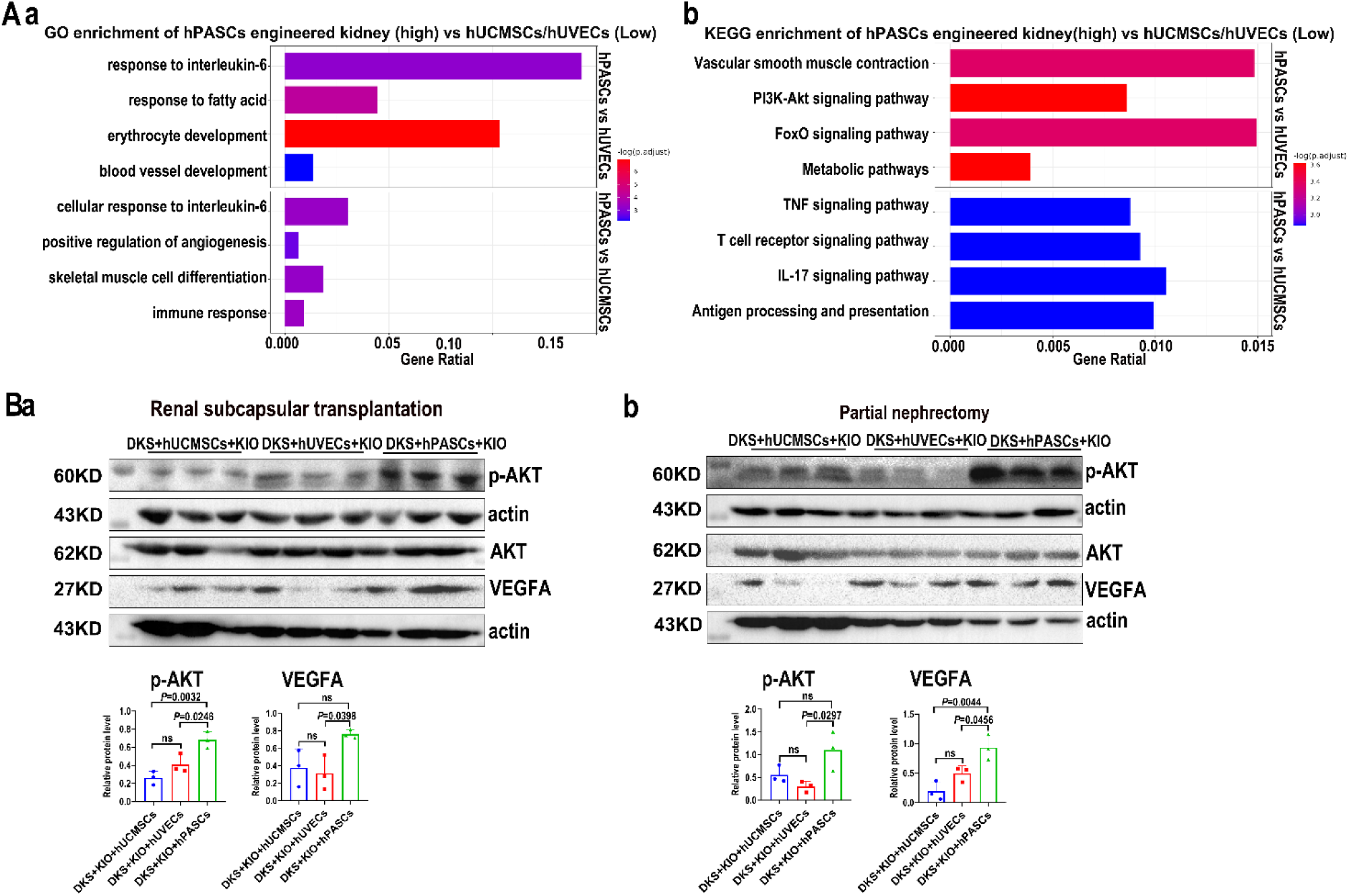
RNA-seq and WB analysis of hPASCs vascularized bioengineered kidney. (A), RNA-seq analysis implants: (a), GO enrichment analysis; (b), KEGG enrichment analysis; *n*=3, *P*<0.05. (B), Western blotting analysis of pAKT and VEGFA protein: (a), implant in renal subcapsular transplantation model; (b), implant in partial nephrectomy and transplantation model: *n*=3, *P*<0.05.

## Notes

### Competing Interest Statement

The authors have declared no competing interest.

